# Glutathione acts as an exometabolite that rescues functionally distinct genetic mutations in fission yeast

**DOI:** 10.64898/2026.01.21.700948

**Authors:** Ryotaro Yoshizumi, Shunichi Miura, Hiroaki Matoba, Go Hirai, Shumpei Asamizu, Hiroyasu Onaka, Yoko Yashiroda, Akihisa Matsuyama, Minoru Yoshida, Shinichi Nishimura

## Abstract

Microorganisms in nature form communities through diverse interactions, such as mutualism and competition, to adapt to their ecological environments. These interactions seem to be mediated by extracellular metabolites (exometabolites), yet the chemical and biological diversity underlying these processes remains largely unexplored. In this study, we examined the chemical basis of cell-cell communication in the fission yeast *Schizosaccharomyces pombe* by a genome-wide screen employing 3,420 viable gene deletion mutants. We identified 37 strains that exhibited growth defects in monoculture on a minimal medium but exhibited growth recovery in the vicinity of wild-type colonies (co-culture), suggesting that exometabolites secreted by wild-type cells compensated for the gene deletion. Both lipophilic and water-soluble fractions obtained by solvent partitioning of the wild-type culture supernatant promoted growth recovery. Among the 11 mutants rescued by the water-soluble fraction, six were cysteine auxotrophs, prompting analyses of thiol-containing metabolites by liquid chromatography-mass spectrometry (LC-MS), revealing the presence of glutathione (GSH) in the culture supernatant. GSH restored growth in most strains as a nutrient source. In contrast, GSH rescued cell morphology defects in the *hob3*Δ mutant, lacking the Bin/Amphiphysin/Rvs (BAR) adaptor protein Hob3, through a mechanism independent of nutrition. This research advances understanding of exometabolite-mediated interactions in *S. pombe* by highlighting the role of GSH as one of the communication molecules that influence cellular processes and shape microbial communities.

**Author summary:** Microorganisms secrete a wide range of metabolites that control microbial community behavior. These extracellular metabolites (exometabolites) include not only well-studied signaling molecules but also diverse primary and secondary metabolites, suggesting complex interactions among microbes. However, the molecular basis of these interactions remains poorly understood, partly due to challenges in detecting them experimentally. In this study, we surveyed exometabolites involved in cell-cell interactions in the model eukaryotic microorganism *Schizosaccharomyces pombe*. *S. pombe* secretes a wide variety of metabolites, including previously reported nitrogen signaling factors (NSFs) and glutathione identified in this work. By analyzing gene deletion mutants that depend on GSH for growth, we provide new insights into how microbes regulate collective behavior by sharing exometabolites.

## Introduction

Microorganisms secrete metabolites into the environment, which influence microbial community behavior (1–4). These extracellular metabolites, called exometabolites, affect microbial populations and their life cycles. Well-known examples include mating pheromones, autoinducers for quorum sensing, and antibiotics produced for microbial competition, which are categorized as secondary metabolites, nonessential for growth of the producing organisms (5–8). However, exometabolites encompass not only secondary metabolites, but also primary metabolites, such as amino acids and organic acids, essential for cellular anabolism and catabolism, including biosynthesis and energy production (9,10).

Indeed, recent studies highlight the importance of extracellular primary metabolites in mediating cell-cell interactions (11–13). For example, metabolite exchange contributes to the acquisition of drug resistance and to lifespan extension in the budding yeast *Saccharomyces cerevisiae* (14–16). Cells that engage in metabolic cooperation exhibit greater metabolite efflux activity because they cooperate metabolically to export drugs more efficiently, making them more tolerant to antimicrobial drugs. Thus, a diverse range of metabolites can function extracellularly, but their exact nature remains unclear, partly due to challenges in detecting exometabolite-mediated interactions (9,17,18). This is particularly true for cell-cell interactions within the same species, where distinguishing donor and recipient cells is difficult.

The fission yeast *Schizosaccharomyces pombe* is a valuable, eukaryotic model organism for investigating fundamental cellular systems, such as cell cycle, cell morphogenesis, chromosome segregation, etc (19). Research using *S. pombe* is supported by genome-wide resources such as gene deletion strain collections and an ORFeome library, as well as curated databases such as Pombase (20–23). However, cell-cell communication in this species remains poorly understood (24,25), except for the mating mediated by peptide pheromones (7,26) and autotoxin (27). Previously, nitrogen signaling factors (NSFs), oxylipins that are produced by *Schizosaccharomyces* species, were identified as exometabolites that relieve nitrogen catabolite repression (NCR) (28,29). NCR suppresses the uptake and metabolism of less preferred nitrogen sources, such as branched-chain amino acids (BCAA: valine, leucine, and isoleucine) in the presence of preferred nitrogen sources such as ammonium chloride or glutamate (Glu) (8,28,30). NSFs consist of two chemically distinct but functionally equivalent molecules, 10(*R*)-acetoxy-8(*Z*)-octadecenoic acid and 10(*R*)-hydroxy-8(*Z*)-octadecenoic acid, the latter of which was used in this study (referred to as NSF). The biological activity of NSFs was visualized by co-culturing a gene deletion mutant and a wild-type strain. A mutant lacking the *eca39* gene (*eca39*Δ), which encodes a BCAA aminotransferase, was unable to grow independently on media containing Glu even in the presence of BCAA. The growth defect of the *eca39*Δ cells was rescued by co-culturing them with wild-type cells that secrete NSFs on the same medium. We refer to such growth recovery of gene deletion mutant cells in co-culture, which does not occur in monoculture, as adaptive growth. These results suggest that identifying novel pairs of mutant and wild-type strains would lead to the discovery of new exometabolites.

Glutathione (GSH) is a tripeptide composed of three amino acids—Glu, cysteine (Cys), and glycine (Gly)—and serves as a nutrient source in *S. pombe* (31–33). GSH also possesses diverse cellular functions, including antioxidant defense and iron homeostasis (34,35). In *S. cerevisiae*, the secretion and uptake of GSH broaden the viable temperature range (13,36). In *S. pombe*, a GSH uptake transporter has been reported, while neither GSH secretion nor transporters responsible for its export have been reported (32,33,37).

In this study, we conducted a genome-wide screen using a viable gene deletion strain collection to identify novel exometabolites that rescue defective phenotypes of particular mutants. A total of 37 mutants exhibited adaptive growth on defined minimal medium (EMM2) in the vicinity of the wild-type cell colonies. Among them NSF rescued the growth of 14 mutants, but not that of the remaining 23. We analyzed culture supernatants of the wild-type strain to identify exometabolites other than NSF and found that GSH compensates for the growth defect of 8 gene deletion mutants. Among them, GSH was utilized as a nutrient source in 7 mutants after degradation into Cys or Glu. However, GSH also rescued the cell growth and morphology in a mutant lacking *hob3* gene, which encodes a Bin/Amphiphysin/Rvs (BAR) adaptor protein, independently of GSH degradation. Since Hob3 senses and induces membrane curvature and is functionally linked to remodeling the actin cytoskeleton (38), GSH appears to play a novel role in regulating the actin cytoskeleton. These results suggest that GSH serves as an extracellular nutrient and possible signaling molecule for actin remodeling in addition to its role as an intracellular redox-controlling molecule.

## Results

### Screening for mutants whose growth is rescued in the vicinity of wild-type colonies

To search for novel exometabolites that regulate cell-cell communication in fission yeast, we designed a screening program to identify gene deletion mutants that exhibit growth defects in EMM2 and growth recovery in the vicinity of wild-type colonies. For the genome-wide screening, we employed a mutant collection consisting of 3,420 haploid gene deletion mutant strains (Fig 1A) (20). Since EMM2 is a defined medium, we expected to easily identify exometabolites secreted by fission yeast. The screening program consists of three steps. First, we compared the growth of mutants on YES, a complex rich medium, and EMM2 to identify mutants that show a growth defect only on EMM2 (S1 Fig). After testing the reproducibility, we identified 77 mutant strains that grew on YES medium but showed poor or no growth on EMM2 (S1 Table). These genes were enriched in amino acid biosynthesis and sulfate assimilation (S2 Table), which is consistent with the previous genome-wide phenotypic profiling of deletion mutants across multiple medium, including EMM2 (39). In the second step, the 77 mutants were inoculated near wild-type colonies on EMM2 agar medium and were co-cultured at 30 °C for 4 days. Of the 77 mutants, 32 showed adaptive growth (Fig 1B). Since 15 of them were involved in mitochondrial respiration, we examined other respiration-related mutant strains that showed poor or no growth both on EMM2 and YES (S1 Fig), resulting in the identification of 7 additional mutant strains that showed adaptive growth (S2 Fig). In the third step, the 39 strains in total were subjected to colony PCR, and the correct gene disruption was confirmed for 38 strains (See Methods). Finally, we examined the reproducibility of the adaptive growth at 27 °C and identified 37 genes whose deletion results in poor growth in monoculture on EMM2 but shows adaptive growth near a wild-type colony (co-culture), probably due to receiving metabolites secreted from the wild-type cells (Table 1, S2 Fig, S3 Table). It is notable that *hob3*Δ and *ipk1*Δ cells exhibited temperature-dependent growth: they failed to grow at 27 °C but grew poorly at 30 °C (S3A Fig).

**Fig 1.**
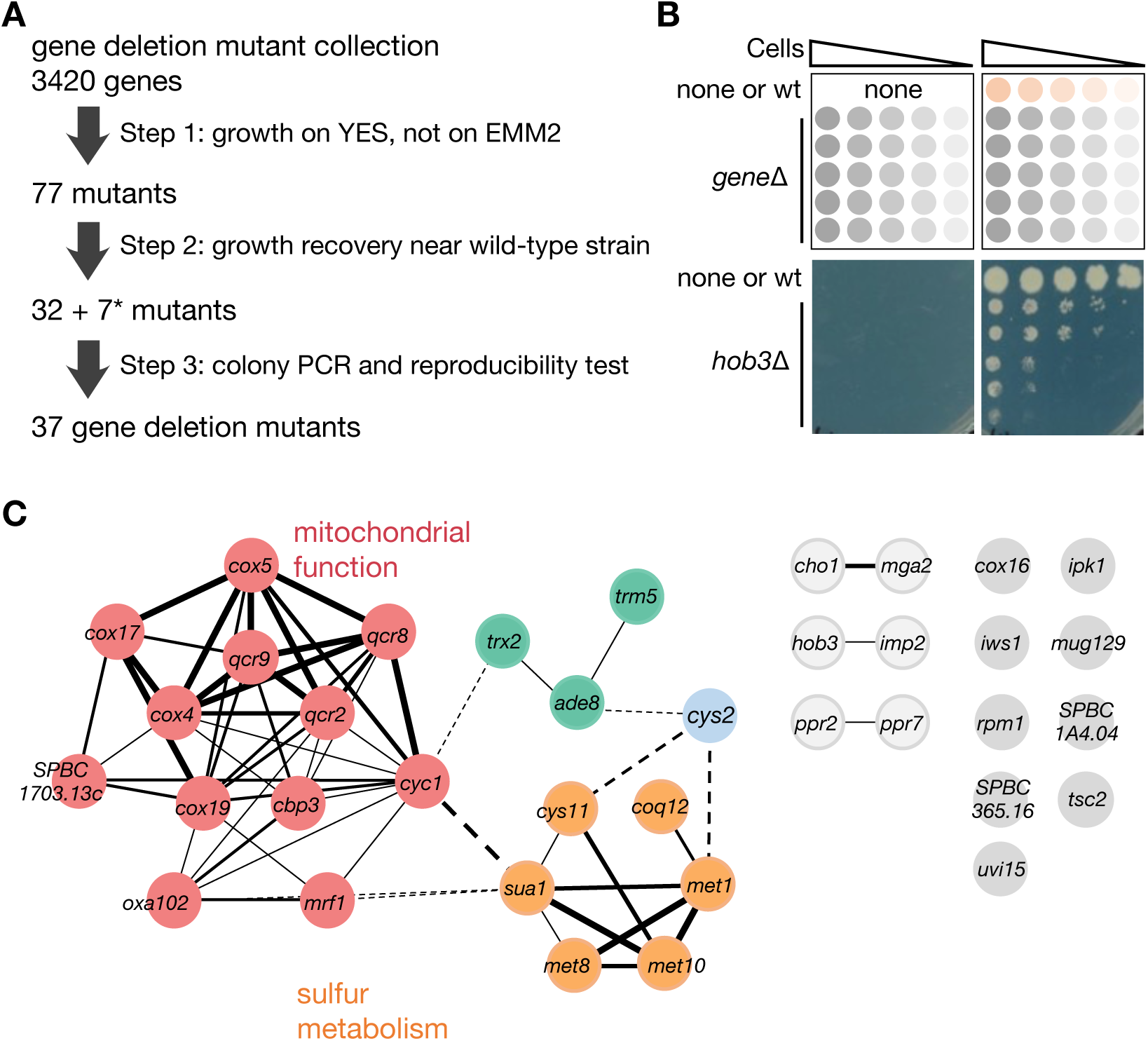
Identification of gene deletion mutants that show adaptive growth. (A) Schematic illustrating the screening process. *Seven mutants involved in electron transport chain were added after the second step. They grew poorly on both YE and EMM2 in the first step. (B) One of the representative screening results. *hob3* Δ exhibited adaptive growth on EMM2 after 7 days of incubation at 27 °C. (C) Network analysis of the 37 genes/proteins using the STRING app. Color coding indicates clusters, and line thickness represents confidence.

**Table 1.**
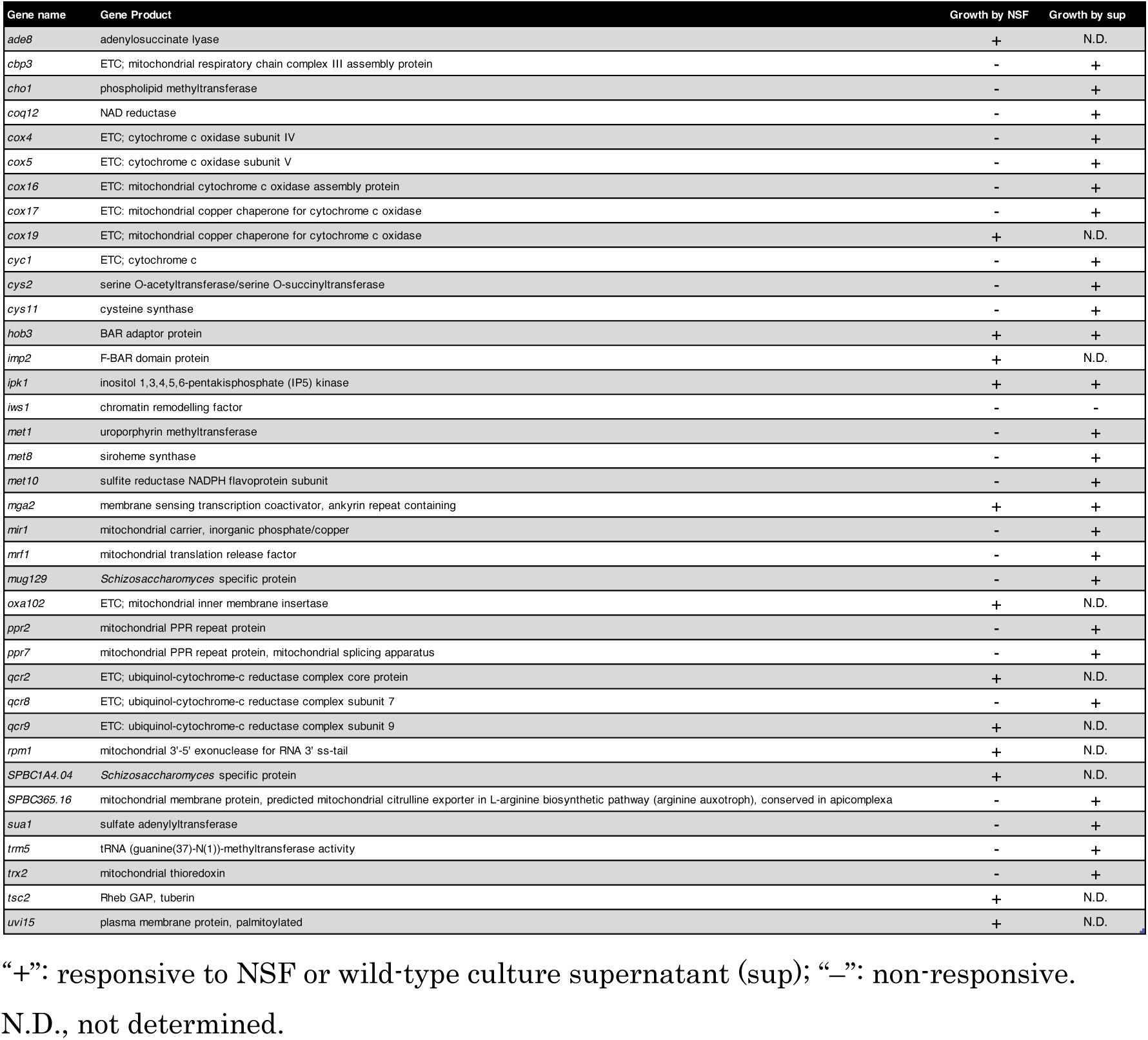
List of 37 gene deletion mutants that show adaptive growth.

Analyses of the Gene Ontology (GO) terms revealed significant enrichment of the 37 genes in mitochondrial ATP synthesis-coupled electron transport (*P* < 1×10^-5^) and sulfur amino acid biosynthetic processes (*P* < 1×10^-3^) (S4 Fig, S4 Table) (40). Network analyses were also carried out using interaction data deposited in BioGRID (Fig 1C) (41). As expected, two clusters of proteins, consisting of mitochondria-associated proteins and sulfur metabolism-associated proteins, were identified. These results suggest that mitochondrial functions and sulfur metabolism can be supported by exometabolites in fission yeast. Furthermore, since half of the genes were not involved in the two clusters, the mechanisms of the adaptive growth seem to be diverse, to which multiple exometabolites may contribute.

### Growth recovery by the culture supernatant

Previously, we identified NSFs as pheromone-like molecules that revoke NCR in *S. pombe* (28,30). Before conducting bioassay-guided fractionation, we tested the effect of NSF on the growth of the 37 mutants at 30 °C (Fig 2A). We observed that 14 mutants exhibited growth recovery in the presence of NSF: *ade8*Δ, *cox19*Δ, *hob3*Δ, *imp2*Δ, *ipk1*Δ, *mga2*Δ, *mrf1*Δ, *oxa102*Δ, *qcr2*Δ, *qcr9*Δ, *rpm1*Δ, *SPBC1A4.04*Δ, *tsc2*Δ, and *uvi15*Δ mutants (Fig 2A, B). Among these, *hob3*Δ and *ipk1*Δ mutants, which showed temperature-dependent growth (S3A Fig), responded to NSF at 30 °C, but not at 27 °C (S3B Fig). Since the wild-type cells rescued their growth at 27 °C, exometabolites other than NSFs may be involved in inducing the adaptive growth of *hob3*Δ and *ipk1*Δ cells, or they may function cooperatively with NSFs.

**Fig 2.**
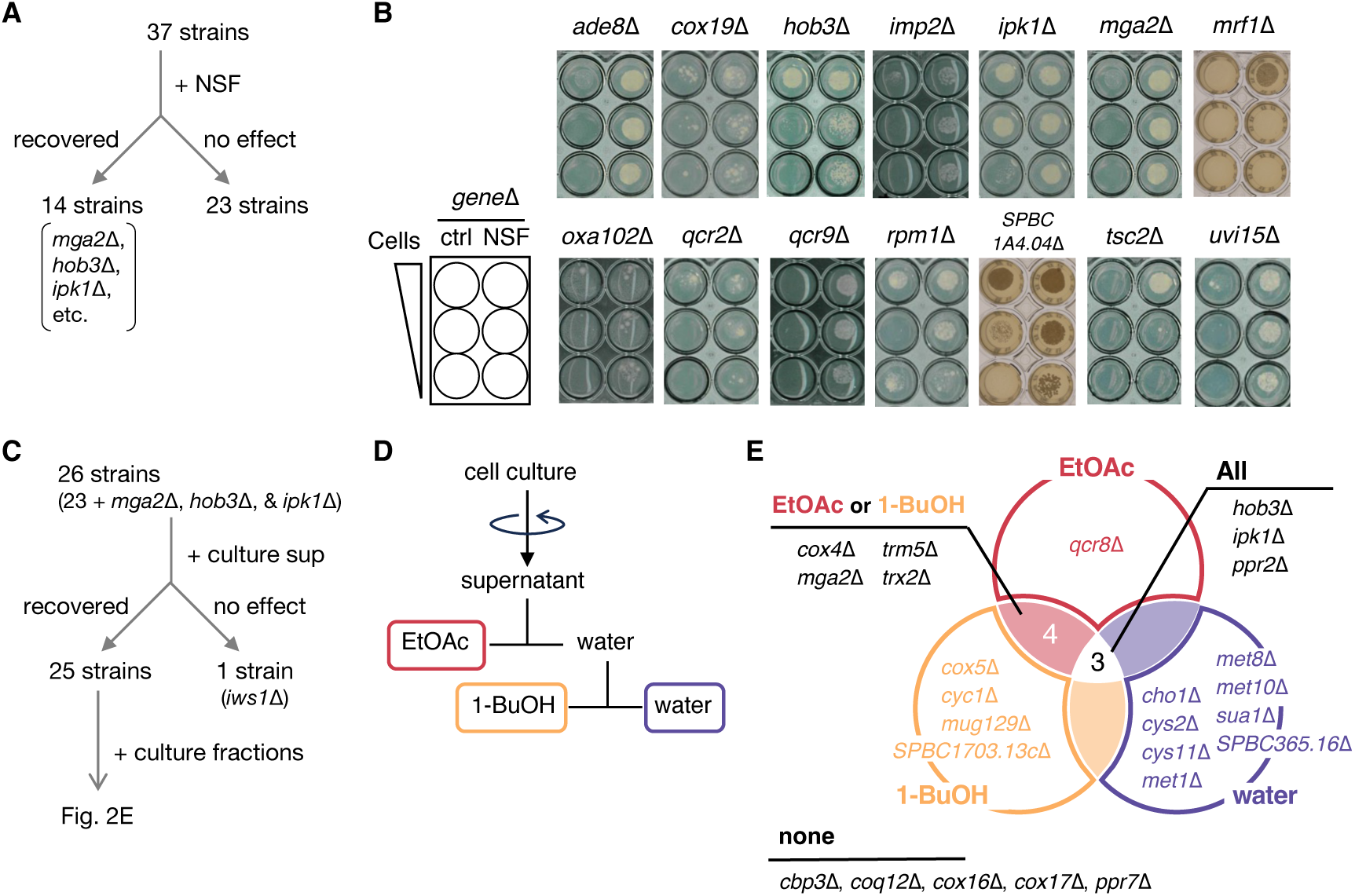
Characterization of 37 strains by their response to NSF or culture supernatants. (A) Summary of the effect of NSF at 30 °C. Among 37 strains tested, 14 displayed growth recovery in the presence of NSF, whereas 23 exhibited no response. (B) Growth recovery of the 14 NSF-responsive strains shown in (A). Serial dilutions of cells were spotted onto media with NSF or 50 % MeOH (ctrl), and growth was assessed visually after 7-14 days at 30 °C. (C) Growth recovery assay using culture supernatants. The 23 NSF-nonresponsive strains and 3 NSF-responsive strains (*mga2*Δ, *hob3*Δ, and *ipk1*Δ) were inoculated with culture supernatant. Among 26 strains, 25 exhibited growth recovery. (D) Fractionation of the culture supernatant. The culture supernatant obtained by centrifugation of wild-type culture was extracted sequentially with ethyl acetate and 1-butanol, to obtain 3 fractions. (E) Growth recovery of the 25 strains (shown in C) by organic and water-soluble fractions. Strains are grouped based on the fractions that induced recovery. Cells were cultured for 7-14 days at 30 °C.

We next examined the activity of the culture supernatant of the wild-type strain to identify novel growth-promoting exometabolites (Fig 2C). For this test, we used 26 disruptants: 23 NSF-nonresponsive strains, one NSF-responsive strain lacking the transcription coactivator (*mga2*Δ) as an NSF-response positive control, and two temperature-sensitive strains (*hob3*Δ and *ipk1*Δ). As expected, the culture supernatant induced growth of most of the mutants, except for *iws1*Δ (S5 Fig). We then sequentially extracted the culture supernatant with ethyl acetate (EtOAc) and 1-butanol (1-BuOH) to obtain three fractions: EtOAc, 1-BuOH, and water-soluble fractions (Fig 2D). The activity of the three fractions was examined using the 25 disruptants (Fig 2C). All three fractions exhibited growth recovery activities, suggesting the presence of multiple active metabolites (Fig 2E, S6 Fig). The *mga2*Δ cells, which responded to NSF, showed growth recovery in the presence of the EtOAc or 1-BuOH fraction. We analyzed these two fractions using gas chromatography-mass spectrometry (GC-MS) and found that NSF was present only in the EtOAc fraction (S7 Fig), suggesting that the 1-BuOH fraction contains active metabolites other than NSF. Five disruptants, including *cbp3*Δ, *coq12*Δ, *cox16*Δ, *cox17*Δ, and *ppr7*Δ, did not respond to any fractions. These mutants may require multiple fractions for their growth recovery, or the active substance may have been lost during the fractionation. Notably, the growth of 11 mutants was recovered by the water-soluble fraction.

### Identification of GSH as a functional exometabolite

Among the 11 mutants that responded to the water-soluble fraction, 6 were found to have mutations in genes involved in Cys biosynthesis: *cys2*Δ, *cys11*Δ, *met1*Δ, *met8*Δ, *met10*Δ, and *sua1*Δ (Fig 3A). Some of them are known to be Cys auxotrophs (32,42,43). We hypothesized that the water-soluble fraction contains Cys or other sulfur-containing exometabolites. To test this possibility, we treated the water-soluble fraction with a thiol-reactive reagent, BODIPY-FL maleimide, and analyzed the fraction for the presence of thiol metabolites by liquid chromatography-mass spectrometry (LC-MS). Unexpectedly, BODIPY-labeled Cys was not detected (S8 Fig). Instead, glutathione (GSH) and H_2_SO_3_ (sulfite) were detected as BODIPY-labeled forms (S8 Fig).

**Fig 3.**
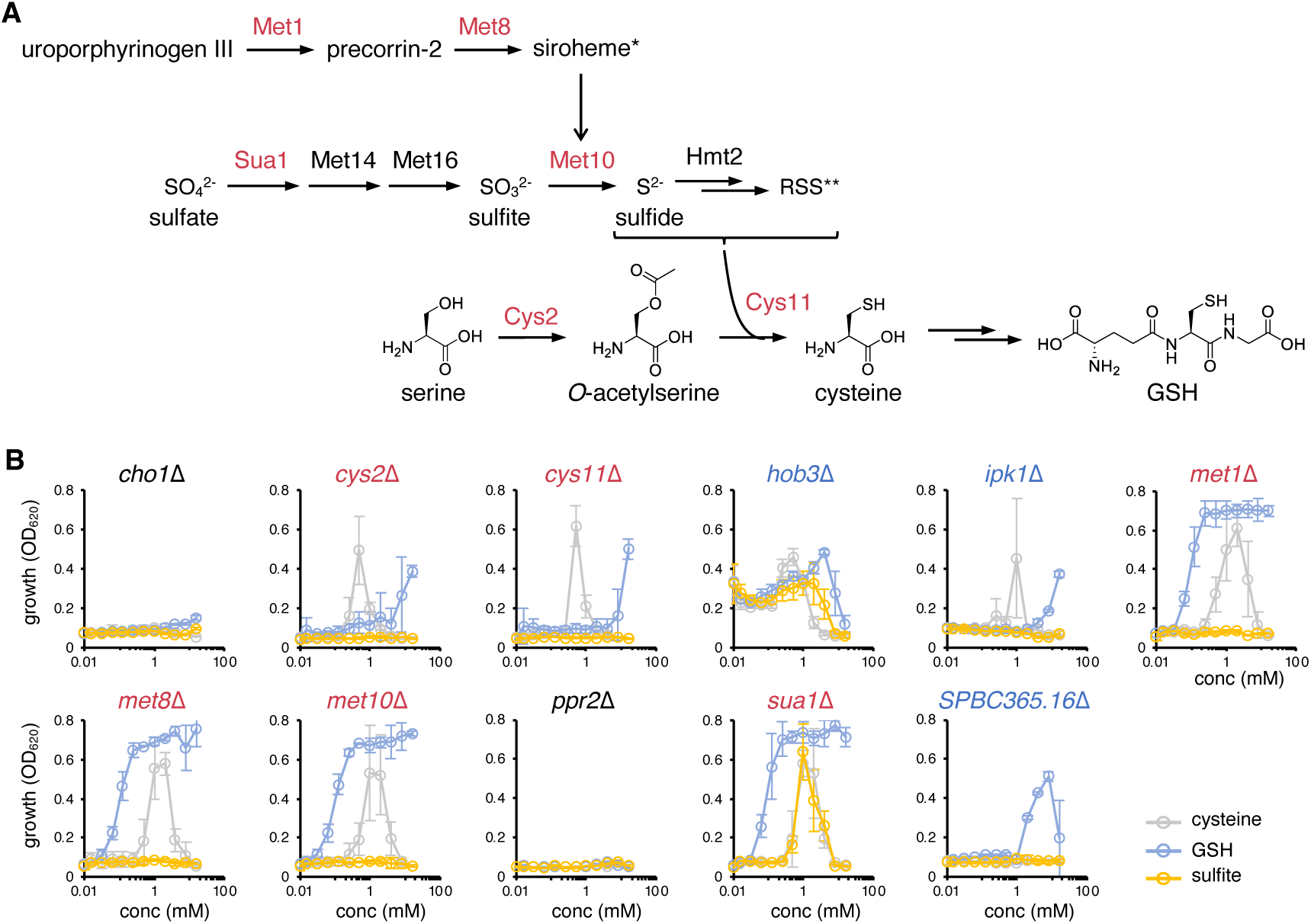
Effect of GSH on the growth of gene deletion mutants. (A) Biosynthetic pathway of cysteine and glutathione (GSH) in *S. pombe*. *Siroheme is required as a coenzyme for sulfite reductase Met10. **Reactive sulfur species (RSS) includes HS_n_H and GSSH. Cys11 can use both H_2_S and GSSH, and probably other RSS. (B) Growth recovery test of the 11 strains that responded to the water-soluble fraction of the culture supernatant. Cells were cultivated with cysteine, GSH, or sulfite for 66 hours at 27 °C. Strains colored in red are Cys biosynthesis deletion mutants (A, B) and those colored in blue are non-Cys biosynthesis deletion mutants that show growth recovery by GSH (B). Data represent the mean ± SD (n = 3).

Next, we conducted growth recovery tests using Cys, GSH, or sulfite in the liquid culture (Fig 3B). The effective concentration was investigated by the serial dilution method. Cys promoted growth recovery in 8 mutants: *cys2*Δ and *cys11*Δ cells at approximately 1 mM; *met1*Δ, *met8*Δ *met10*Δ and *sua1*Δ cells in the range of 1 to 5 mM; *hob3*Δ and *ipk1*Δ cells at approximately 1 mM, although these two mutants did not show recovery on agar media (see below). GSH was also effective on these Cys-responsive strains, albeit at different concentrations: *cys2*Δ and *cys11*Δ cells showed growth recovery by GSH at concentrations higher than 5 mM. In the case of *met1*Δ, *met8*Δ, *met10*Δ, and *sua1*Δ cells, the minimal effective concentration of GSH was lower (100 µM) than that of Cys, and high concentrations did not suppress the growth, indicating that GSH is more efficiently incorporated and metabolized in these mutants. *hob3*Δ and *ipk1*Δ cells also showed growth recovery in response to GSH, although they required slightly higher concentrations of GSH than Cys. The growth of *SPBC365*.*16*Δ cells was recovered only by GSH at concentrations above 2 mM. Sulfite was found to induce growth recovery in *sua1*Δ cells at concentrations around 1 mM, similar to the effect of Cys. *cho1*Δ and *ppr2*Δ cells did not respond to sulfur-containing exometabolites, but *cho1*Δ cells showed growth recovery in the presence of choline chloride, as reported (S9 Fig) (44). Thus, the water-soluble fraction may contain choline or its derivatives.

As the adaptive growth was originally observed on agar media (Fig 1A, B), we tested the effects of Cys and GSH on agar media using 9 mutants that showed growth recovery in the liquid media with Cys and/or GSH (Fig 3B, S10 Fig). Similar to the results obtained in liquid media, the growth of *met1*Δ, *met8*Δ, *met10*Δ, and *sua1*Δ cells was recovered by supplementation with Cys or GSH, and *SPBC365.16*Δ cells recovered growth only in response to GSH. However, unlike the results obtained in liquid media, *cys2*Δ, *cys11*Δ, and *hob3*Δ cells did not respond to Cys, and *ipk1*Δ responded to neither Cys nor GSH on agar media. Although the reason is currently unknown, they may not be able to take up enough nutrients for colonization on solid media. We confirmed that the wild-type colony also releases GSH on the EMM2 agar medium (S11 Fig). Therefore, we concluded that GSH is an exometabolite responsible for inducing adaptive growth in 8 mutants but not in *ipk1*Δ.

### GSH serves as a source of Cys and Glu

To investigate the mechanism underlying the growth recovery by GSH, we focused on the GSH catabolic pathway, because GSH is a tripeptide including Cys and the 6 Cys biosynthesis gene deletion mutants appeared to utilize GSH as a source of Cys. Cys is obtained from GSH by first hydrolyzing the *γ*-glutamyl bond of GSH, followed by cleaving the peptide bond of Cys-Gly by dipeptidyl peptidase or dipeptidase (S12 Fig) (31,45). Alternatively, GSH can be oxidized by Hmt2 to GSSH, which plays an essential role as a sulfite donor for synthesizing Cys by Cys11 (Fig 3A, S12 Fig) (46,47). To identify key enzymes in these pathways, we evaluated the growth of 7 gene deletion mutants: 5 genes encoding putative dipeptidyl peptidase or dipeptidase (Dug1, Dpe1, Dpe2, SPCC757.05c, and Xpa1), one gene encoding the oxidoreductase Hmt2, and another gene encoding the GSH uptake transporter Pgt1 as a positive control (Table 2, S13 Fig) (20). *SPCC757.05c*Δ, *dpe1*Δ, *dpe2*Δ, and *xpa1*Δ cells, as well as the wild-type cells, were able to grow when more than 100 µM GSH was added as a sole sulfur source. On the other hand, *dug1*Δ and *hmt2*Δ cells required higher concentrations of GSH (700 µM) to grow, suggesting that Dug1 and Hmt2 are required to generate organic sulfur from GSH (Fig 4A, S13 Fig). The *dug1*Δ *hmt2*Δ double mutant required a much higher concentration of GSH (1.5 mM) for growth than that for either single mutant, indicating that Dug1 and Hmt2 function in independent GSH degradation pathways. Specifically, a dipeptidase Dug1 likely digests GSH. Note that the *hmt2*Δ single mutant showed no growth when sulfate was used as the sole sulfur source, indicating that Hmt2 is required for assimilating inorganic sulfur, which uses GSH as a cofactor. Consistent with the role of Pgt1 in GSH uptake, *pgt1*Δ cells did not grow when GSH was supplemented at 1 mM or less.

**Fig 4.**
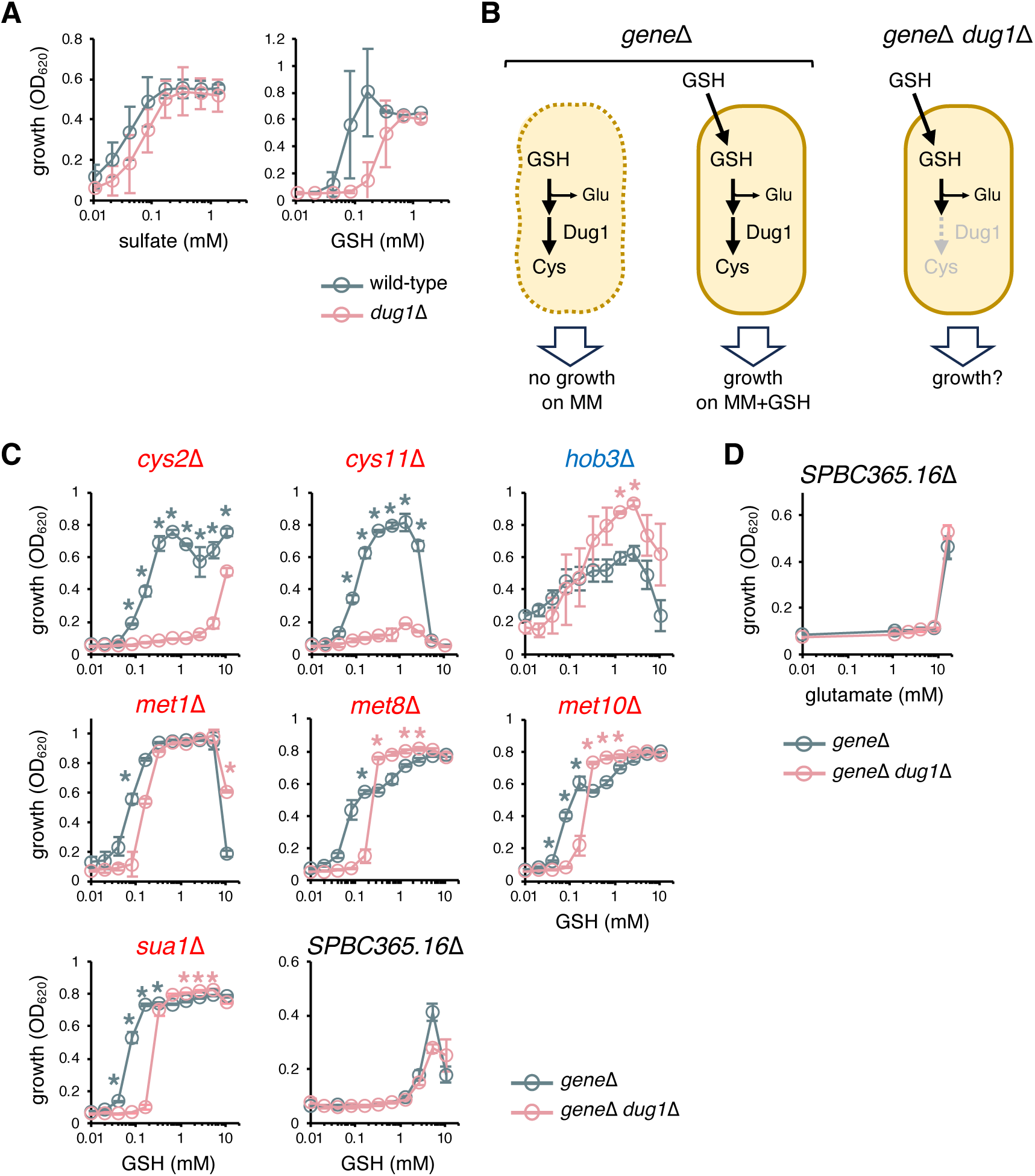
Involvement of Dug1 in GSH-dependent growth. (A) Assimilation of sulfate or GSH as a sulfur source in wild-type and *dug1*Δ strains. Cells were cultivated for 48 hours at 30 °C in EMM2–S medium supplemented with the indicated concentrations of sulfate or GSH. (B) Schematic model illustrating the involvement of *dug1* in GSH assimilation. If a *gene*Δ strain utilizes GSH as a Cys source, growth recovery of the double mutant *gene*Δ *dug1*Δ by GSH would be abolished or attenuated. If GSH is not utilized as an amino-acid source, the double mutant should still show growth recovery by GSH. (C) Effect of *dug1* deletion on the growth recovery by GSH in 8 gene deletion strains. Cells were cultivated with GSH for 66 hours at 27 °C. Six strains labeled in red are Cys biosynthesis deletion mutants. *hob3*Δ labeled in blue showed enhanced growth recovery by *dug1* gene deletion at concentrations 1.3 and 2.6 mM (\**p* < 0.05). *P*-values were obtained from comparisons between *gene*Δ and *gene*Δ *dug1*Δ using Welch’s t-tests and adjusted for multiple comparisons using the Bonferroni correction. * *p* < 0.05. (D) Growth of *SPBC365.16*Δ and *SPBC365.16*Δ *dug1*Δ strains in EMM2 supplemented with Glu for 66 hours at 27 °C. *dug1* deletion had no effect on the Glu utilization. (A, C, D) Data represent the mean ± SD (n = 3).

**Table 2.** Genes tested for the involvement of the GSH assimilation.

| Gene name | Gene/proteine information | Growth test with sulfur source |  |
| --- | --- | --- | --- |
|  | Gene product | Sulfate | GSH |
| *dpe1* | dipeptidyl peptidase, unknown specificity, implicated in glutathione metabolism | growth | growth |
| *dpe2* | dipeptidyl peptidase, unknown specificity, implicated in glutathione metabolism | growth | growth |
| *dug1* | dipeptidase | growth | decreased |
| *hmt2* | sulfide-quinone oxidoreductase | no growth | decreased |
| *pgt1* | plasma membrane glutathione transmembrane transporter (positive control) | growth | no growth |
| *SPBC757.05c* | peptidase family M20 protein involved in glutathione catabolism | growth | growth |
| *xpa1* | X-Pro dipeptidase | growth | growth |

Next, we examined the growth recovery of double gene deletion mutants (*gene*Δ *dug1*Δ), as Dug1 was shown to be involved in the GSH degradation and assimilation (Fig 4A). We hypothesized that deletion of *dug1* would increase the minimal concentration of GSH required for growth recovery, if the mutant relies on GSH as an amino acid source (Fig 4B, 4C). As expected, the minimal effective concentrations increased by deleting *dug1* in the Cys biosynthesis gene deletion mutants. However, GSH still induced growth recovery in *met1*Δ *dug1*Δ, *met8*Δ *dug1*Δ, *met10*Δ *dug1*Δ, and *sua1*Δ *dug1*Δ cells at the concentrations above 100 µM, suggesting that Hmt2 still functions in these mutants. *SPBC365.16*Δ cells did not show a change in the effective concentration upon *dug1* deletion. This gene encodes a mitochondrial membrane protein and deletion mutant was previously shown to grow in the presence of arginine on EMM2 agar media (48), suggesting that GSH is used as a Glu source, a precursor of arginine. Cleavage of Glu from GSH does not require Dug1 (S12 Fig). Indeed, *SPBC365.16*Δ cells exhibited growth recovery in the presence of Glu (Fig 4D), confirming that GSH functions not only as a Cys source but also as a Glu source.

### GSH maintains the cell polarity and robust growth in *hob3*Δ

Unlike other gene deletion strains identified in this study, the *hob3*Δ strain is unique in that its growth was recovered by GSH but not by Cys or Glu on the agar media (Fig 5A). Consistently, the *dug1* deletion in the *hob3*Δ mutant did not reduce the growth recovery but rather enhanced the effect of GSH on growth recovery (Fig 4C, Fig 5A). To confirm that GSH functions after incorporation into cells, we generated a *hob3*Δ *pgt1*Δ strain because Pgt1 is the sole GSH transporter in *S. pombe* (32). In this strain, the growth recovery by GSH was abolished (Fig 5A). These results indicate that GSH itself functions in cells to promote *hob3*Δ growth recovery, rather than through its metabolic products.

**Fig 5.**
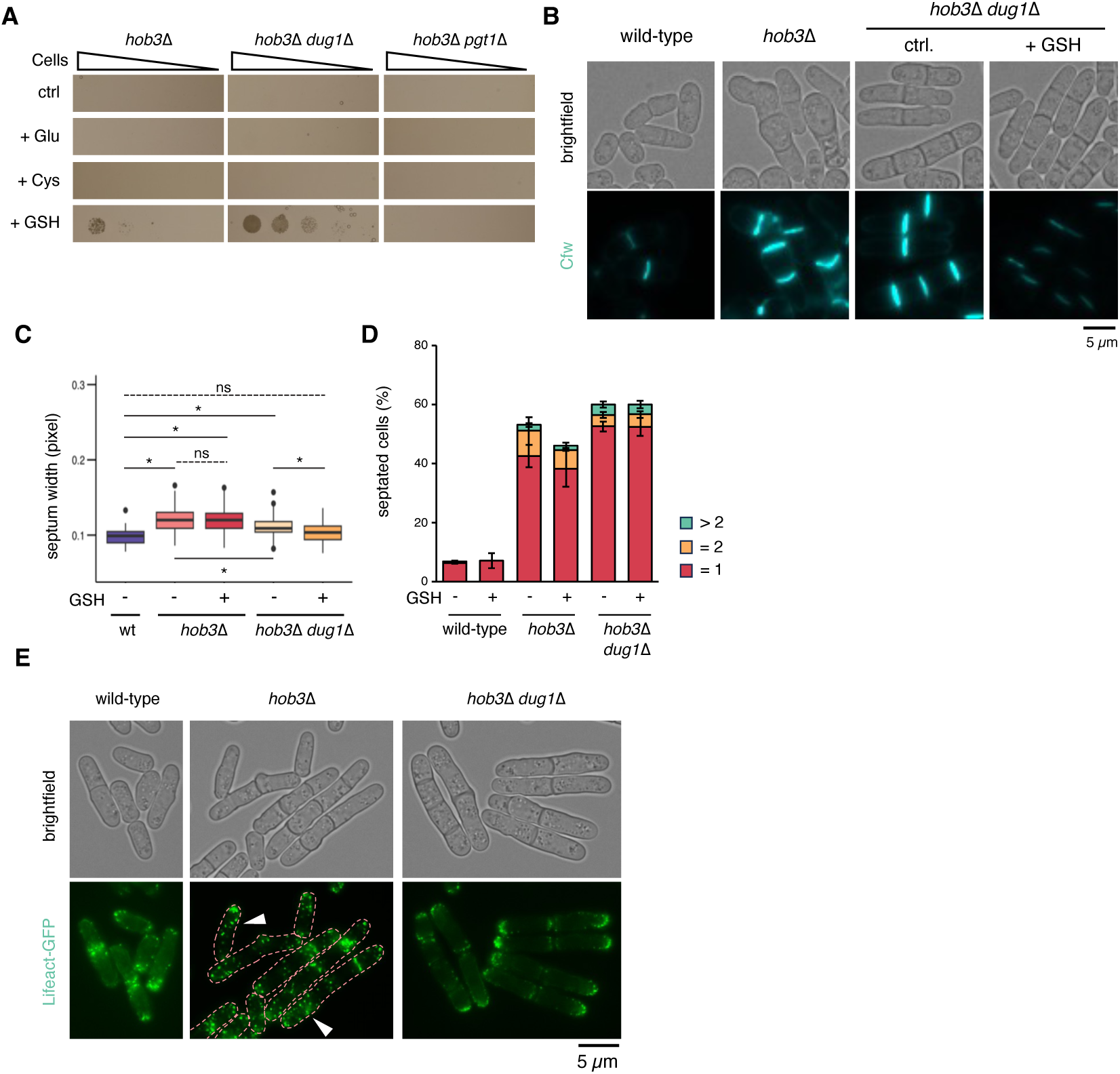
GSH maintains cell polarity and contributes to the growth recovery of *hob3*Δ. (A) Growth of *hob3*Δ, *hob3*Δ *dug1*Δ, and *hob3*Δ *pgt1*Δ cells on EMM2 (top), EMM2 supplemented with glutamate, cysteine, or GSH (from top to bottom). Cells were cultured for 10 days at 27 °C. (B) Effect of GSH on cell morphology and septa (stained by Cfw) of wild-type, *hob3*Δ, and *hob3*Δ *dug1*Δ cells. Cells were cultured in EMM2 with or without 1 mM GSH at 30 °C and observed by fluorescence microscopy. Scale bar = 5 µm. (C) Cell width at the septum in wild-type, *hob3*Δ, and *hob3*Δ *dug1*Δ cells with or without GSH at 30 °C. *P*-values were determined using one-way ANOVA with Bonferroni’s multiple comparison test. ns, not significant. * *p* < 0.0001. (D) Population of cells with one, two, or more septa in wild-type, *hob3*Δ, *hob3*Δ *dug1*Δ strains at 30 °C. Data represent the mean ± SD (n = 3). (E) F-actin distribution visualized by lifeact-GFP in wild-type, *hob3*Δ, and *hob3*Δ *dug1*Δ cells at 30 °C. White arrows indicate broader distribution of F-actin patches in *hob3*Δ cells. Scale bar = 5 µm.

Hob3 is a BAR adaptor protein that senses and induces membrane curvature. It is also functionally linked to remodeling the actin cytoskeleton (38). In *S. pombe*, the *hob3* deletion leads to multiseptated cells with intense calcofluor white (Cfw) staining, which labels β-(1,3)-glucan in the cell wall (49), and mislocalized F-actin, likely due to improper localization of Cdc42 to the cell division sites (50,51). To examine the effect of GSH, we analyzed cell morphology. As previously reported, *hob3*Δ cells exhibited a rounded cell shape with an intense fluorescent signal of Cfw (Fig 5B, S14 Fig). The cell width at the septum in *hob3*Δ cells was wider than that in wild-type cells (Fig 5C). Although GSH treatment did not affect the cell width of *hob3*Δ cells, deleting *dug1* decreased it. Treating *hob3*Δ *dug1*Δ cells with GSH further decreased the cell width, making it comparable to that of wild-type cells. Another feature of *hob3*Δ cells is the increased population of septated cells: the number of cells with a septum drastically increased when compared with the wild-type cells, and multiseptated cells appeared (Fig 5D). Most *hob3*Δ cells had curved septa. GSH treatment did not reduce the population of septated *hob3*Δ cells, while the curved septa were still found in GSH-treated *hob3*Δ cells (S14 Fig). Importantly, we could not observe the curved septa in *hob3*Δ *dug1*Δ cells, although the Cfw fluorescent signal remained intense (Fig 5B, S14 Fig). This intense Cfw signal was almost completely attenuated by GSH treatment in *hob3*Δ *dug1*Δ cells.

To investigate the F-actin structure, we expressed lifeact-GFP, a probe that specifically binds to F-actin, in the mutant cells (52,53). Although the dumpy cell shape of *hob3*Δ cells was somewhat alleviated by expressing lifeact-GFP, which can potentially influence the actin dynamics, a difference in the F-actin patch distribution was observed between *hob3*Δ and *hob3*Δ *dug1*Δ cells (Fig 5E) (54). F-actin patches were mainly detected in the cell tip and around the septum in the wild-type cells. In contrast, they were randomly dispersed in *hob3*Δ cells. This abnormality was rescued by deleting *dug1* (Fig 5E). When expressing lifeact-GFP, by which dumpy cell shape of *hob3*Δ cells were corrected, exogenous GSH did not show detectable changes in the cell morphology including actin patch distribution in either *hob3*Δ cells or *hob3*Δ *dug1*Δ cells (S15 Fig). This contrasts with the GSH-dependent improvements in septum morphology and Cfw staining (Fig 5B, 5C), and with the effect of *dug1* deletion on the lifeact-GFP localization (Fig 5D). These results suggest that the effect of GSH on the abnormal cell morphology of *hob3*Δ is exerted through correcting F-actin structures, which can be masked by expressing lifeact-GFP.

The *hob3*Δ cells grew slowly in EMM2 liquid medium at 30 °C, with a growth rate approximately half that of the wild-type cells (S16 Fig). Notably, the *dug1* deletion in the *hob3*Δ mutant restored the growth rate to the level of the wild-type cells. Supplementation with GSH did not restore the growth rate of *hob3*Δ cells but promoted the growth rate of both *hob3*Δ *dug1*Δ and *dug1*Δ cells at 30 °C. Collectively, these data indicate that: (i) *dug1* deletion partly normalizes growth (S16 Fig), septum shape (Fig 5B), cell width (Fig 5C), and F-actin localization (Fig 5E) in *hob3*Δ, and (ii) exogenous GSH further improves growth (Fig 5A, S16 Fig), intense Cfw signal (Fig 5B) and cell width (Fig 5C) in the *dug1*Δ background. These findings suggest that the growth defects in *hob3*Δ cells can be remedied by inhibiting the degradation of GSH and increasing cellular GSH levels, likely through regulation of cell morphology.

## Discussion

Our functional genomic screen identified 37 gene deletion strains that showed adaptive growth when co-cultured with the wild-type strain on EMM2. These 37 genes were required for growth in monoculture on EMM2, and the growth defect of these mutants appeared to be compensated by exometabolites secreted by the wild-type colonies. GSH was identified as one of the exometabolites promoting adaptive growth in 8 mutants. Seven strains utilized GSH as a Cys or Glu source, while GSH restored the growth of *hob3*Δ through a different mechanism. Abnormal septum shape and mislocalization of actin patches in *hob3*Δ cells were restored by deleting the *dug1* gene and supplementing with GSH.

GSH consists of three amino acids, and therefore serves as an amino acid source (31). In our study, 6 mutants lacking genes involved in the Cys biosynthesis pathway displayed growth recovery in the presence of GSH: *cys2* Δ, *cys11*Δ, *met1*Δ, *met8*Δ, *met10*Δ, and *sua1*Δ cells (Fig 3B, 4C, S10 Fig). These mutants used GSH as a Cys source. This notion was supported by the result that the effective concentration of GSH increased upon deletion of *dug1*, encoding a putative dipeptidase that cleaves the dipeptide Cys-Gly into Cys and Gly (Fig 4C). In most strains, the minimal effective concentration of GSH was lower, and its effective concentration range was broader than that of Cys (Fig 3B). At high concentrations, Cys was toxic, whereas GSH was not. GSH also served as a Glu source: growth of *SPBC365.16*Δ cells was recovered by Glu, as well as GSH (Fig 4C, D). The minimal concentration of Glu required for growth recovery was higher than that of GSH. These results suggest that extracellular GSH is a more effective, safer amino acid source than Cys or Glu under our experimental conditions.

There are two pathways for obtaining Cys from GSH: the Dug1-dependent pathway and the Cys11-dependent pathway (S12 Fig) (45,46). This is supported by the finding that *cys11*Δ *dug1*Δ cells barely assimilated GSH as a Cys source (Fig. 4C). Notably, deleting *dug1* in *cys2*Δ cells resulted in a 64-fold increase in the minimal GSH concentration for growth recovery. In *met1*Δ, *met8*Δ, *met10*Δ, and *sua1*Δ cells, however, deleting *dug1* increased effective GSH concentrations only by 2- to 4-fold. Redundant enzymes and pathways may exist for Cys2, which acts just upstream of Cys11, and for Met1, Met8, Sua1, and Met10, which are further upstream of Cys11 (Fig 3A). *cys2*Δ and *cys11*Δ cells showed growth recovery by GSH, but the effective concentration of GSH was about 100-fold higher than that of the other 4 mutants (*met1*Δ, *met8*Δ, *met10*Δ, and *sua1*Δ cells) (Fig 3B), which again suggests the presence of redundant enzymes/pathways for these upstream mutants.

NSF, a known oxylipin that regulates NCR, also promoted adaptive growth in *hob3*Δ cells. However, this growth recovery was temperature-dependent: *hob3*Δ showed adaptive growth in response to NSF at 30 °C but not at 27 °C (Fig 2B, S3 Fig). Since adaptive growth of *hob3*Δ cells near wild-type colonies was observed at 27 °C, the presence of exometabolites other than NSF was suggested. Indeed, GSH was identified as the responsible metabolite. Multi-septation and disorder of actin patches were observed in *hob3*Δ cells, which were drastically restored by deleting *dug1* and supplementation with GSH simultaneously (Fig 5). This suggests that GSH may be depleted in *hob3*Δ cells. Although the underlying mechanism is unknown, decreased production, increased degradation, decreased uptake, and/or increased consumption of GSH might occur in *hob3*Δ cells. GSH treatment alone was not sufficient to restore the morphological phenotype in *hob3*Δ cells. Since Hob3 controls the subcellular localization of Cdc42, a key signaling protein that regulates membrane trafficking (51,55), deletion of *hob3* may impair the localization of the GSH transporter Pgt1, resulting in reduced GSH uptake.

There are several possible explanations for how GSH restores the *hob3*Δ phenotypes. One possibility is the elimination of reactive oxygen species (ROS). An actin mutant strain produces elevated levels of ROS, and deleting *hob3* may similarly lead to excess ROS production (56). Extracellular GSH might scavenge the increased ROS (34,36,57). Another possibility is the post-translational modification of proteins by GSH, such as actin (58,59). In budding yeast, actin molecules are stabilized when thiol groups of Cys residues are protected with *N*-acetylcysteine (60). The same mechanism may apply in fission yeast. Finally, GSH can activate the sulfur metabolic pathway. GSH is a substrate for Hmt2 and can therefore activate the Cys11-dependent metabolic pathway (46). Recently, NSF was shown to interact with and activate Hmt2 to revoke NCR (61). In the present study, we found that NSF promoted adaptive growth in *hob3*Δ cells (Fig 2B). Therefore, GSH and NSF may share a common molecular mechanism. However, these possibilities are not mutually exclusive, and the involvement of other unexpected mechanisms should also be considered, which makes elucidating the relationship between *hob3*Δ cells and GSH a compelling challenge.

NSFs revoke NCR in *eca39*Δ and *leu1-32* cells that cannot synthesize BCAAs (28). Growth recovery of *leu1-32* cells by NSFs requires the leucine transporter Agp3. NSF physically interacts with Hmt2, thereby activating respiration and consequently revoking NCR (61). In this study, 14 gene deletion mutants displayed growth recovery by NSF (Fig 2B), and plausible mechanisms can be inferred for some of them. Some mutants likely have defects in mitochondrial functions, e.g. Qcr2 and Qcr9, which function in the electron transport chain. It is reasonable to assume that mitochondrial dysfunction represses the activity of Hmt2, which is relieved by direct activation of Hmt2 by NSF. NSF restored the growth of *tsc2*Δ cells, possibly by activating leucine uptake via Agp3, since *tsc2*Δ cells are known to be defective in leucine uptake (62). *mga2*Δ cells were also rescued by NSF: it is possible that NSF synthesis is reduced in the *mga2*Δ cells, because Mga2 is a transcription factor regulating lipid metabolism. Therefore, the synthesis of oleic acid, the presumed biosynthetic precursor of NSFs (28), is likely reduced in this mutant (63). Alternatively, NSF is incorporated into phospholipids as a substitute for oleic acid in the mutant. Other mutants have no predictable mechanism of action based on current knowledge. Elucidating the mode of action of NSF in the 14 mutants will lead to a deeper understanding of the mode of action of NSFs and the functions of these gene products.

The fission yeast *S. pombe* cells likely secrete multiple exometabolites that support growth, in addition to NSFs and GSH. The water-soluble fraction restored growth in 11 mutants (Fig 2E). While 8 mutants responded to GSH, the other three did not: *cho1*Δ, *ppr2*Δ, and *ipk1*Δ cells (Fig 3B, S10 Fig). Cho1 catalyzes the methylation of phosphatidylethanolamine in the biosynthesis pathway of phosphatidylcholine (PC) (44). The growth defect of the *cho1*Δ cells was rescued by choline (Fig 9). This suggests that the water-soluble fraction may contain choline or related metabolites. Since methylated compounds like choline mediate interactions between marine bacteria and their host microalgae ̶enabling bacteria to escape the lag phase by sensing metabolically active host—, it is intriguing to investigate the physiological role of extracellular choline or methylated metabolites in *S. pombe* (64). The 1-BuOH fraction, which did not contain NSFs, restored growth defect in 11 mutants, suggesting the presence of novel exometabolites involved in cell-cell communications (Fig 2E, S7 Fig).

Microorganisms adapt to a variety of environmental conditions, such as nutrient limitation, extreme temperatures, and fluctuating osmotic pressure. The mechanisms by which they adapt remain poorly understood, but they sometimes achieve this through sharing exometabolites. In this study, we screened *S. pombe* gene deletion mutants on minimal medium and identified GSH as one of the exometabolites that supports the robust growth of this organism. The effective use of mutant collections will enable systematic analyses of cell-cell communication within a species and reveal novel exometabolites and their physiological roles.

## Materials and methods

### Yeast general methods

Strains and oligonucleotide primers used in this study are listed in S5 and S6 Table. Correct gene disruption in the Bioneer yeast deletion library v5.0 was verified by colony PCR using primer pairs kan0/cp5 and kan5/cp3 (20). For *arg5*Δ, whose deletion was not verified in the third step, both the up-tag and down-tag sequences were amplified by colony PCR and sequenced, revealing that the strain was *trx2*Δ. Strains constructed in this study were generated either by genetic crossing or via a PCR-based gene deletion strategy (65,66). In the latter case, *dug1*Δ and *pgt1*Δ were generated by replacing the coding regions with the hygromycin B phosphotransferase gene using the pHph-GFP plasmid, which was constructed by replacing the kanMX marker of pFA6a-kanMX-GFP with the hygromycin resistance (hph) marker from pAG32 (67,68).

Cells were cultured in YES medium or Edinburgh Minimal Medium 2 (EMM2) (69). Depending on auxotrophic requirements, 75 mg/L Ade, 37.5 mg/L Ura, and 75 mg/L Leu were added to EMM2. To prepare EMM2–S medium, sodium sulfate was omitted, and sulfate-containing minerals were replaced with sulfate-free alternatives (Fig 4, S13 Fig). SPA medium was used for genetic crossing when generating double deletion mutants (70).

Cys, sodium hydrogen sulfite, GSH, and choline chloride were added to media at the indicated concentrations. NSF was synthesized as previously described (28). For bioassay, NSF was dissolved in 50% methanol (MeOH) at 1.25 µg/mL, and 20 µL of the solution was applied onto 48-well agar plates and allowed to dry (Fig 2B). Lyophilized culture supernatant was processed in the same manner (S5 Fig, S6 Fig).

### Screen for mutants that show adaptive growth

In the first step, deletion strains from the Bioneer collection were cultured in YES liquid medium at 30 °C for 2 days, then transferred onto either YES or EMM2 agar plates using a 96-pin replicator. Plates were incubated at 30 °C for 4 days. Colony growth was assessed by autofluorescence using the Odyssey imaging system (LI-COR) with an excitation wavelength of 685 nm and emission detection at 700 nm. In the second step, 77 candidate strains identified in the first step were spotted adjacent to wild-type colonies on agar plates and incubated at 30 °C for 6 days. Growth enhancement was assessed visually. In the third step, among the 39 candidate mutant strains, correct gene deletion in 38 was verified by colony PCR. These 38 mutants were then spotted next to wild-type colonies and incubated at 27 °C, identifying 37 strains that exhibited adaptive growth reproducibly.

### Network analysis

Genetic and protein–protein interaction data were retrieved from the STRING (https://string-db.org/) and BioGRID (https://thebiogrid.org/) databases for genes whose deletion exhibited adaptive growth (40,41). Interaction networks obtained from BioGRID were visualized using Cytoscape.

### Preparation of culture supernatant

Wild-type cells (JY1) were pre-cultured in YES liquid medium at 27 °C overnight. Cells in the exponential growth phase were washed twice with EMM2 medium, inoculated into fresh EMM2 liquid medium, and cultured at 27 °C for 48 hours. The cultures were then centrifuged to separate the supernatant and cell pellet. The supernatant was lyophilized and dissolved in 50% MeOH to obtain concentrated samples for the growth recovery assay.

For fractionation, an equal volume of ethyl acetate was added to the culture supernatant, and the mixture was vortexed and centrifuged to separate the fractions. This extraction step was repeated twice, and the organic fractions were combined and designated as the ethyl acetate fraction. Then the aqueous fraction was extracted with an equal volume of 1-butanol, twice. The combined supernatant was designated as the 1-BuOH fraction. The ethyl acetate, 1-buthanol, and water-soluble fractions were concentrated using a rotary evaporator or lyophilizer. All fractions were dissolved in 50% MeOH to make 10-times concentrated solution.

### Detection of NSF in the EtOAc fraction

To the lyophilized sample of 2 mL of culture supernatant, 30 µL of pentadecanoic acid solution (10 µg/mL) as an internal standard and 2 mL of ultrapure water were added. Lipophilic molecules were extracted with an equal volume of ethyl acetate twice, concentrated, lyophilized, and dissolved in 30 µL of *N*-Methyl-*N*-trimethylsilyl trifluoroacetamide (MSTFA, Wako). The mixture was vortexed for 1 minute and incubated at 70 °C for 15 minutes. Derivatized samples were analyzed using a GCMS-QP2010 SE (Shimadzu) equipped with a DB-23 column (0.25 mm × 60 m, 0.15 µm film thickness; Agilent Technologies). Each sample was injected in split mode (1:50) at 80 °C. After an initial isothermal hold at 50 °C for 1 minute, the column temperature was increased at a rate of 25 °C/min to 175 °C, then at 4 °C/min to 230 °C. The carrier gas was helium at a flow rate of 0.66 mL/min. The derivatized NSF showed characteristic peaks at m/z 299.00 and 329.15 (28).

### Detection of thiol compounds in the aqueous fraction

The water-soluble fraction of culture supernatants or water-soluble extracts of agar media were lyophilized and derivatized as follows. To each culture sample, 1 M Tris-HCl (pH 8.0; final concentration 100 mM), BODIPY-FL maleimide (final concentration 8 µg/mL, BroadPharm), and Milli-Q water were added. The mixtures were vortexed, incubated at 30 °C for 12 hours in the dark, and subjected to HPLC coupled with photodiode array (PDA) or electrospray ionization mass spectrometry (ESI-MS) in both positive and negative ion modes. HPLC was performed on a Chromolith® Performance RP-18e column (100 mm × 4.6 mm, Merck) at a flow rate of 0.8 mL/min. A linear gradient from 5% to 55% acetonitrile over 22 minutes was followed by 100% acetonitrile for additional 2 minutes. Detection was carried out by absorption at 503 nm using photodiode array (PDA) detector. In high-resolution MS analysis (negative mode), Peak 1 showed a main ion at *m/z* 720.2451, with accompanying isotopic peaks at 719.2473 and 721.2474. Peak 2 showed a main ion at m/z 495.1334, with corresponding signals at 494.1363 and 496.1355. These spectra were consistent with the predicted BODIPY adducts of GSH (C₃₀H₃₈BF₂N₇O₉S) and sulfite (C₂₀H₂₃BF₂N₄O₆S), respectively. Standard compounds were prepared by reacting commercially available GSH or sodium sulfite with BODIPY-FL maleimide. The identity of the detected peaks was confirmed by comparison of the sample spectra with those of the standards in both HPLC retention time and ESI-MS spectral patterns.

### Fluorescence microscopy

Cells in the exponential growth phase were incubated in EMM2 medium with or without 1 mM GSH at 30 °C for 16 hours. Calcofluor white (Sigma-Aldrich) was added to the culture at a final concentration of 1 µg/mL using a 1 mg/mL aqueous stock solution (71). Images were acquired using a KEYENCE BZ-X710 fluorescence microscope equipped with a PlanApo 60× objective and BZ-X Viewer software. Cell width was measured using the straight-line tool in ImageJ, based on pixel length.

### Statistical analyses

In Fig 4C, Statistical analyses were performed to compare growth between *gene*Δ and *gene*Δ *dug1*Δ strains for each concentration within the same strain background, and *P*-values were calculated using Welch’s t-tests and adjusted for multiple comparisons using the Bonferroni correction. In Fig 5B, *P*-values were calculated using one-way ANOVA followed by Bonferroni’s multiple comparison test. In S15 Fig, multiple testing correction was applied using the Benjamini–Hochberg method (FDR).

## Supporting information

## Acknowledgements

We are grateful to the members of the Laboratory of Microbiology, the Chemical Genomics Research Group, Molecular Ligand Target Research Team, and Drug Discovery Seed Compounds Exploratory Unit for technical support and helpful discussions. English editing was partly done with the aid of ChatGPT and Copilot.

## Author contributions

Conceptualization: RY SM AM MY SN.

Formal analysis: RY SM SN.

Funding acquisition: AM YY MY SN.

Investigation: RY SM (a part of screen) SA(MS) HO (MS) AM (plasmid construction).

Methodology: RY SM AM SA HO MY SN.

Project administration: RY SM AM MY SN.

Supervision: AM MY SN.

Writing – original draft: RY SN.

Writing – review & editing: AM YY MY SN.

