## Supplementary Figures for "Glutathione acts as an exometabolite that rescues functionally distinct genetic mutations in fission yeast"

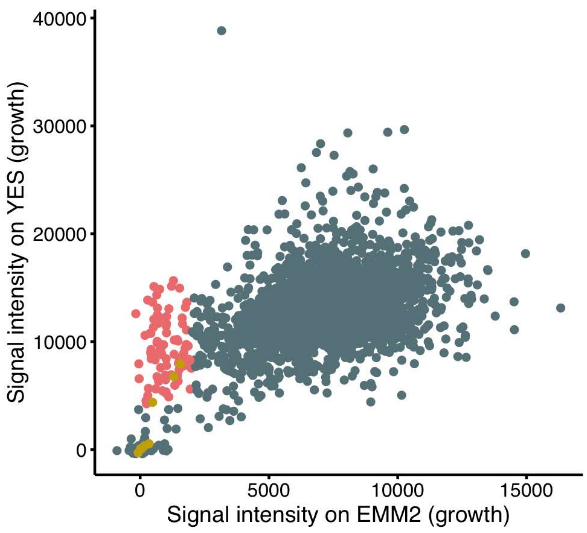


**S1 Figure. Comparison of growth of gene deletion mutants on rich and minimal media.** Results of the growth test from the first step of the screen are shown. Signal intensities of the mutant colonies on EMM2 or YES medium are plotted. Mutants with signal intensities above 4,000 on YES and below 2,000 on EMM2 (red) were selected for the second step. Additional respiration-related mutant strains with signal intensities below 2,000 on EMM2 (yellow) were included for further analyses.

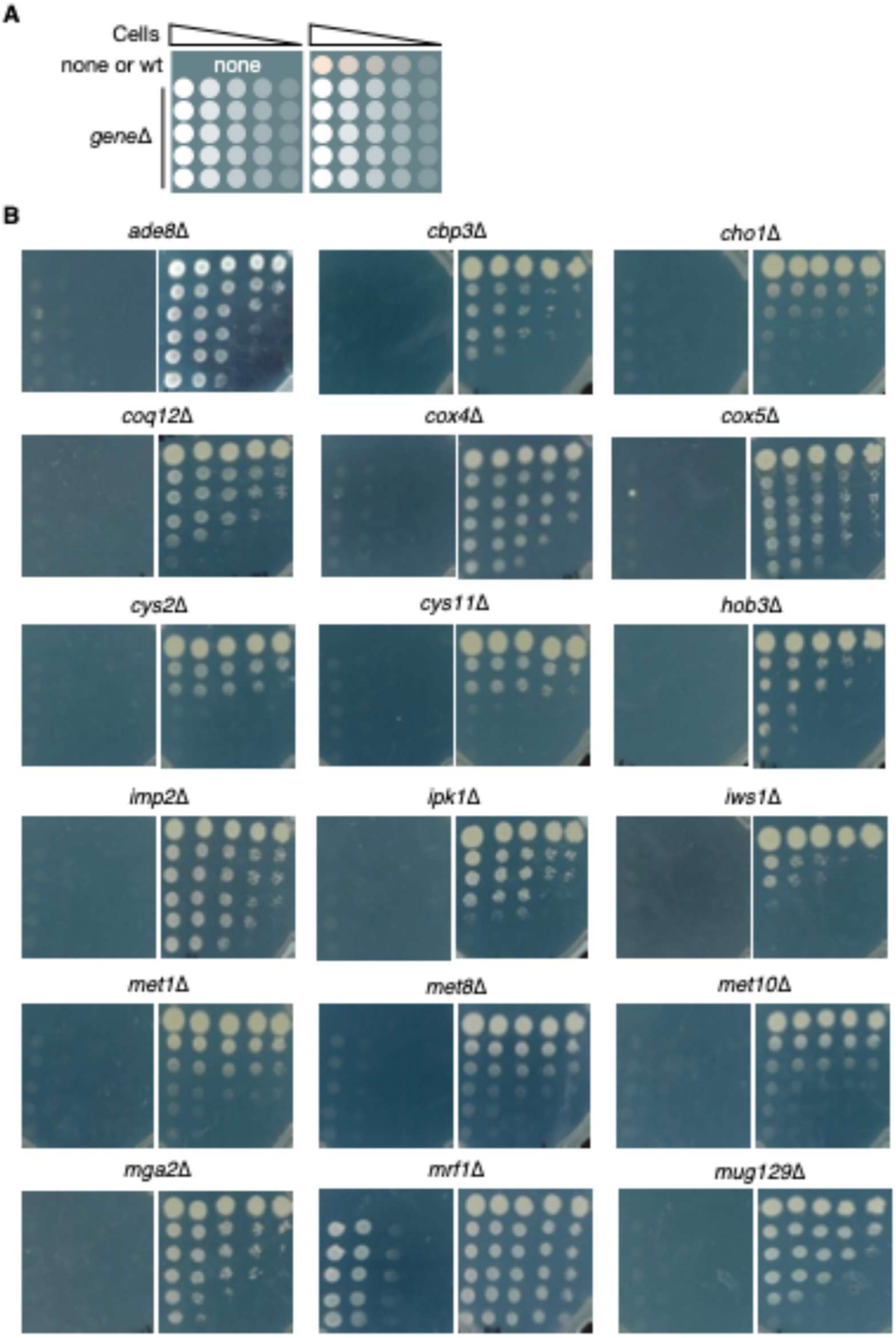


**S2 Figure. Adaptive growth of 37 strains identified at 3 steps of the screen.**

(A) Schematic illustration of the growth assay. Serial dilutions of the mutant cells were spotted onto EMM2 with or without co-culture with the wild-type strain.

(B) Adaptive growth of 30 strains identified in the third step, among the 32 strains selected in the second step. For the *arg5*Δ strain, gene deletion was not verified by PCR, and, in fact, the strain was *trx2*Δ. *hal4*Δ cells did not exhibit adaptive growth in the third step. Serial dilutions of cells were spotted onto EMM2 and incubated at 27 ˚C for 7 to 14 days.

(C) Adaptive growth of 7 mitochondrial gene deletion strains, among the 8 more respiration-related mutant strains added in the second step. They grew poorly on both YES and EMM2 media in the first step. *cox6*Δ cells did not exhibit adaptive growth in the second step. Serial dilutions of cells were spotted onto EMM2 and incubated at 27 ˚C for 7 to 14 days.

Representative images are shown (n = 2–3).

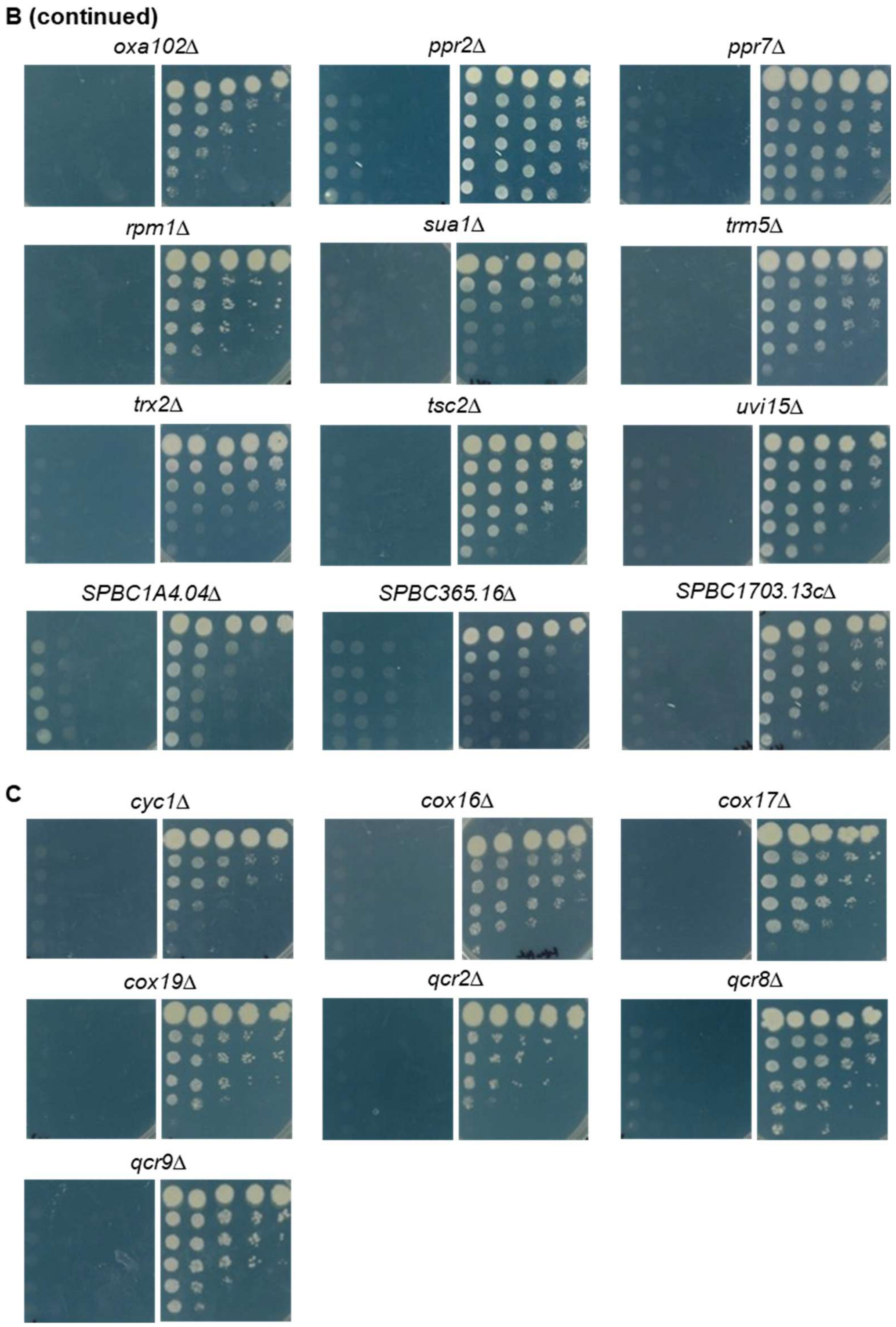


S2 Figure (continued)

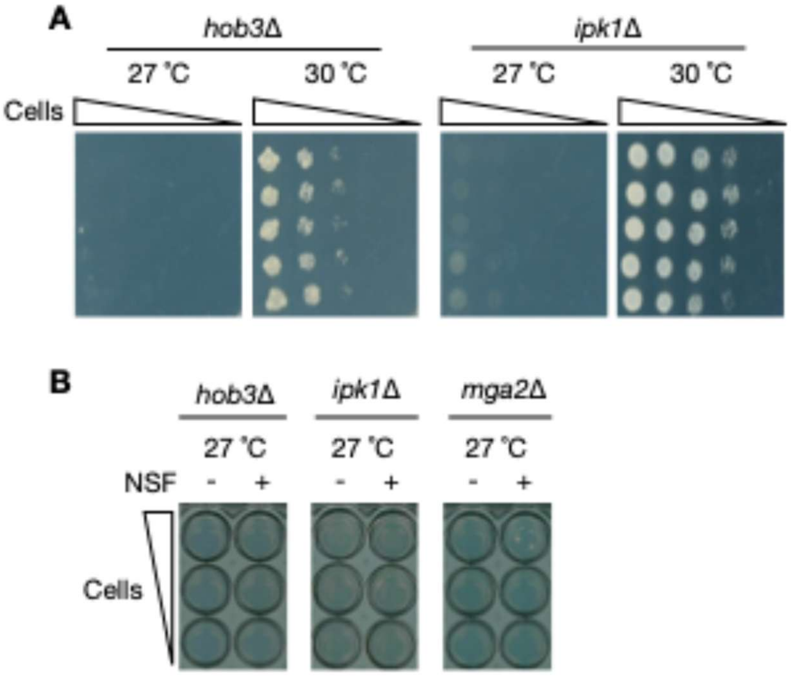


**S3 Figure. Temperature-dependent growth of *hob3*Δ and *ipk1*Δ strains.**

(A) Temperature-dependent growth of *hob3*Δ and *ipk1*Δ strains. Serial dilutions of cells were spotted onto EMM2, and incubated at 27 ˚C or 30 ˚C.

(B) *hob3*Δ and *ipk1*Δ did not respond to NSF at 27 ˚C, whereas *mga2*Δ did at 27 ˚C.

Representative images of 2 independent experiments are shown.

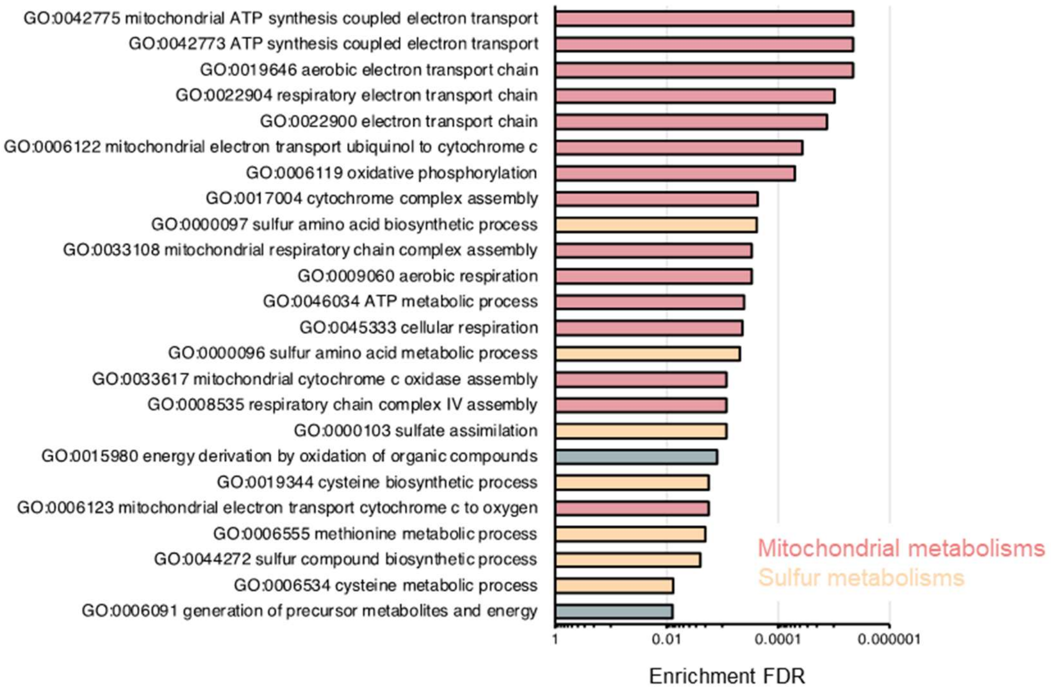


**S4 Figure.** GO enrichment analysis of biological processes was performed for the 37 strains that showed adaptive growth using Shiny GO 0.80. GO terms with $p < 0.01$ are shown.

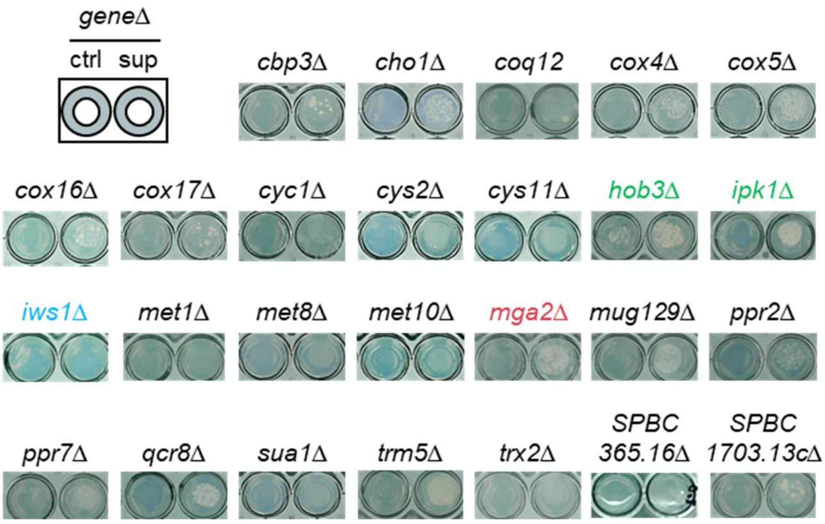


**S5 Figure. Growth recovery by the culture supernatant of the wild-type strain.**

Mutant cells were spotted onto EMM2 supplemented with either 50% methanol (ctrl) or the culture supernatant of the wild-type strain (sup) and incubated at 30 °C. Growth was assessed visually. Strains colored in green are temperature-sensitive mutants (*hob3*Δ and *ipk1*Δ), the strain colored in red is an NSF-responsive strain (*mga2*Δ, a positive control), and the strain colored in blue (*iws1*Δ) is the only one that did not respond to the culture supernatant. Representative images of 2-3 independent experiments are shown.

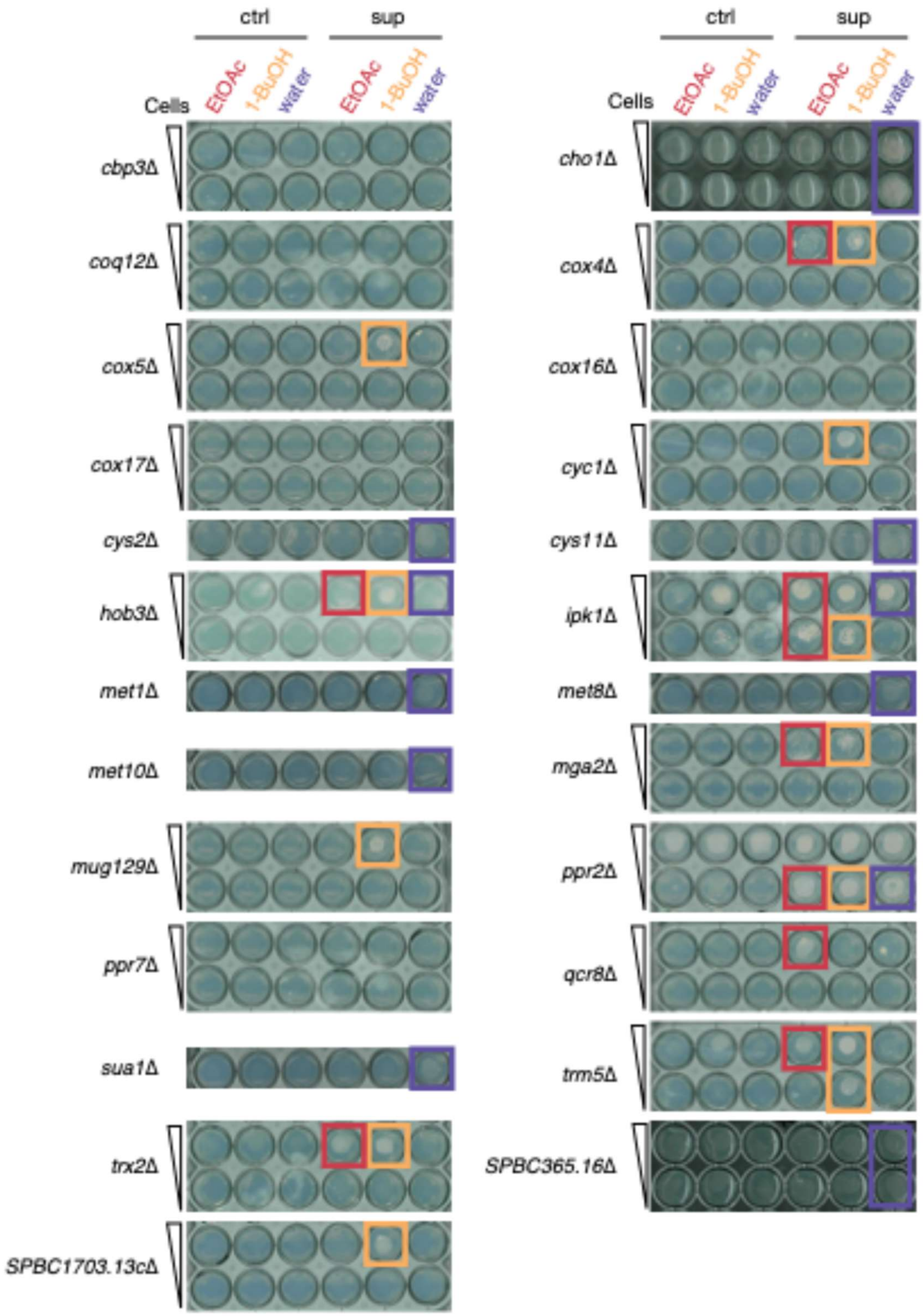


**S6 Figure. Growth recovery by crude fractions derived from culture supernatants.**

Mutant cells were spotted onto EMM2 supplemented with either fractions derived from EMM2 (ctrl) or from the culture supernatant of the wild-type strain (sup) and incubated at 30 °C. Growth confirmed visually is indicated with colored square. For some strains, serial dilutions were spotted. Representative images of 2–3 independent experiments are shown.

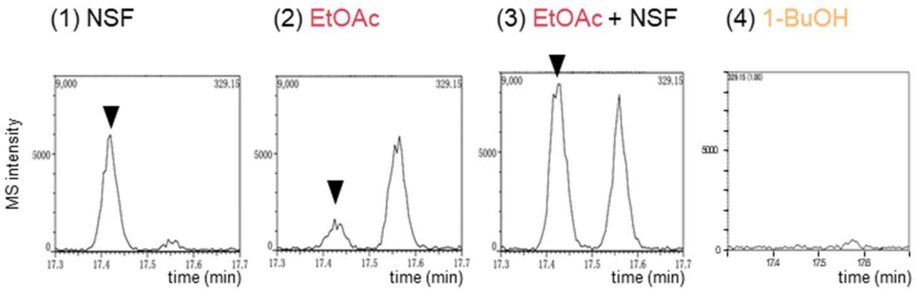


**S7 Figure. Detection of NSF in the EtOAc extract of the culture supernatant.**

Extracted ion chromatograms (EIC) at *m/z* 329.15 in the GC-MS analyses are shown. Synthetic 10(*R*)-hydroxy-8(*Z*)-octadecenoic acid (NSF) (1), the EtOAc fraction of the culture supernatant (2), a mixture of NSF and the EtOAc fraction (3), and the 1-BuOH fraction (4) were subjected to GC-MS analyses. Peaks corresponding to NSF are indicated by black arrow heads. The co-injection analyses (3) unambiguously showed that NSF was eluted at 17.4-17.45 min. All samples were trimethylsilylated prior to GC-MS analysis. Representative images in 3 independent experiments are shown.

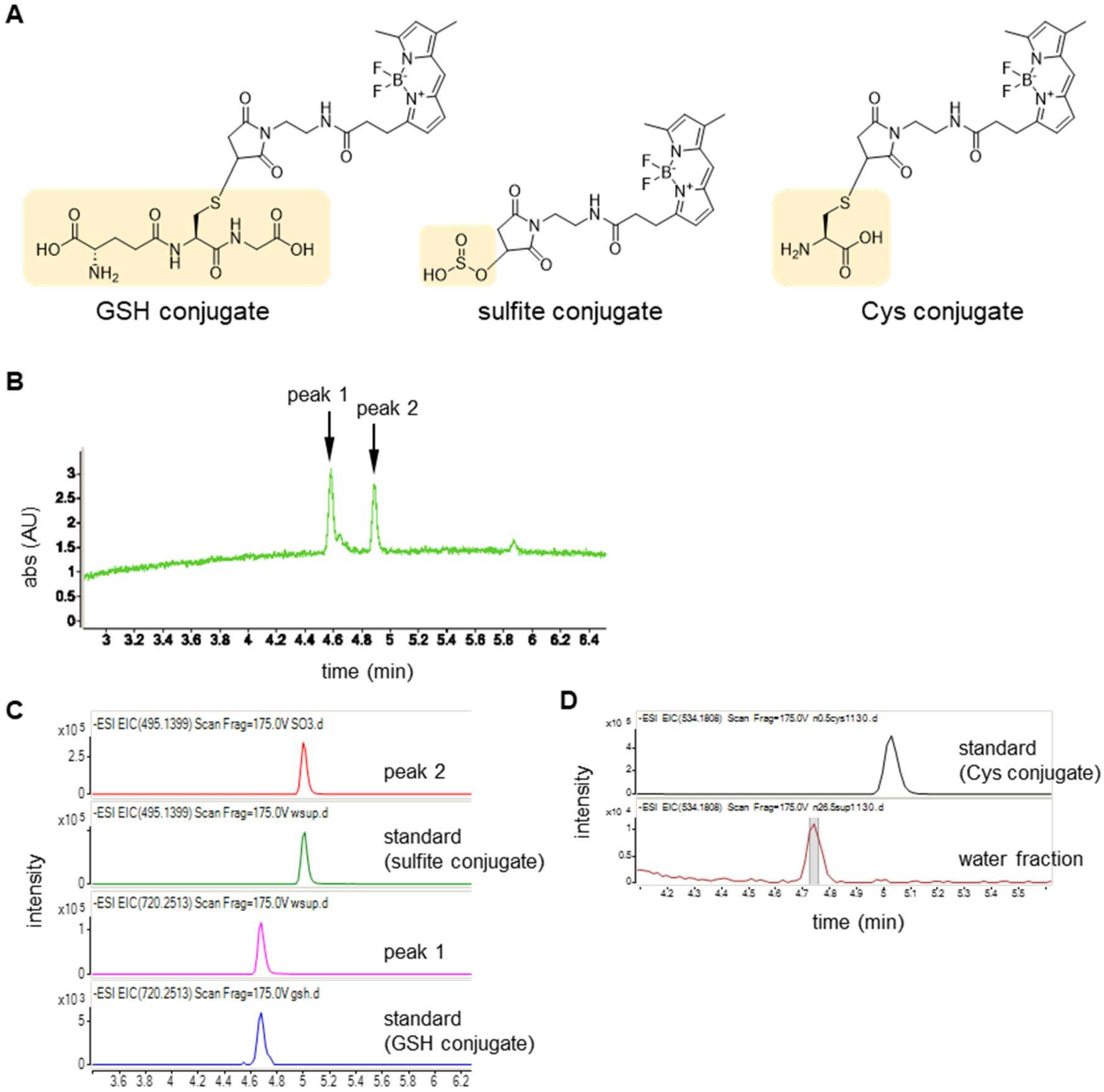


**S8 Figure. Detection of GSH and sulfite in the water fraction.**

(A) Chemical structures of BODIPY-labeled GSH, sulfite, and Cys.

(B) Photodiode array (PDA) analysis of the water fraction at 503 nm. Two peaks were detected.

(C) EICs at *m/z* 495.1399 ± 0.2476 and 720.2513 ± 0.2671. The retention time of peak 1 matched that of the sulfite conjugate, while the retention time of peak 2 matched that of the GSH conjugate.

(D) EIC at *m/z* 534.1808 ± 0.3601. No peak corresponding to the Cys conjugate was detected in the water fraction.

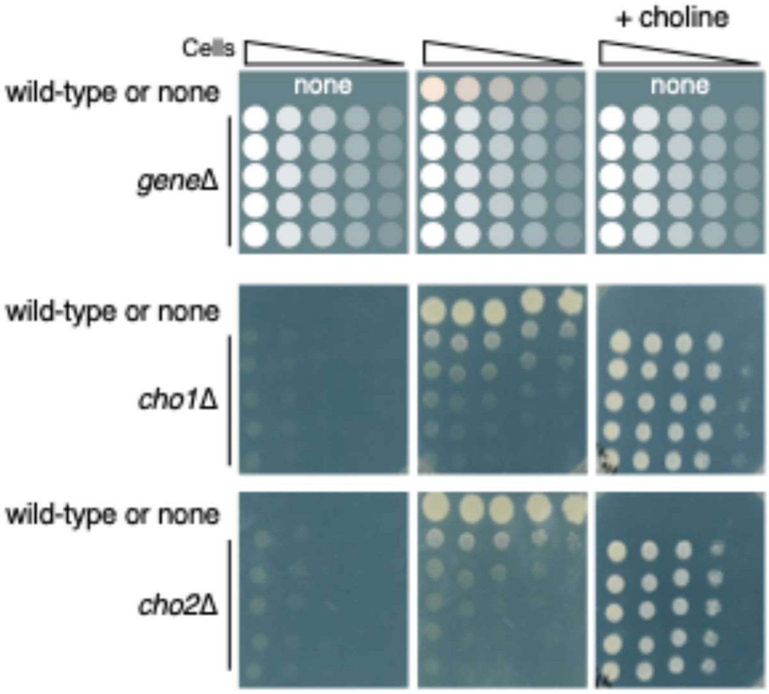


**S9 Figure. Growth recovery of *cho1*Δ and *cho2*Δ by wild-type colony and choline.**
Serially diluted *cho1*Δ or *cho2*Δ cells were spotted onto EMM2 alone (left), EMM2 with wild-type cells (center), or EMM2 supplemented with 100 μM choline chloride (right) and incubated at 27 °C for 11 days. Representative images in 2 independent experiments are shown.

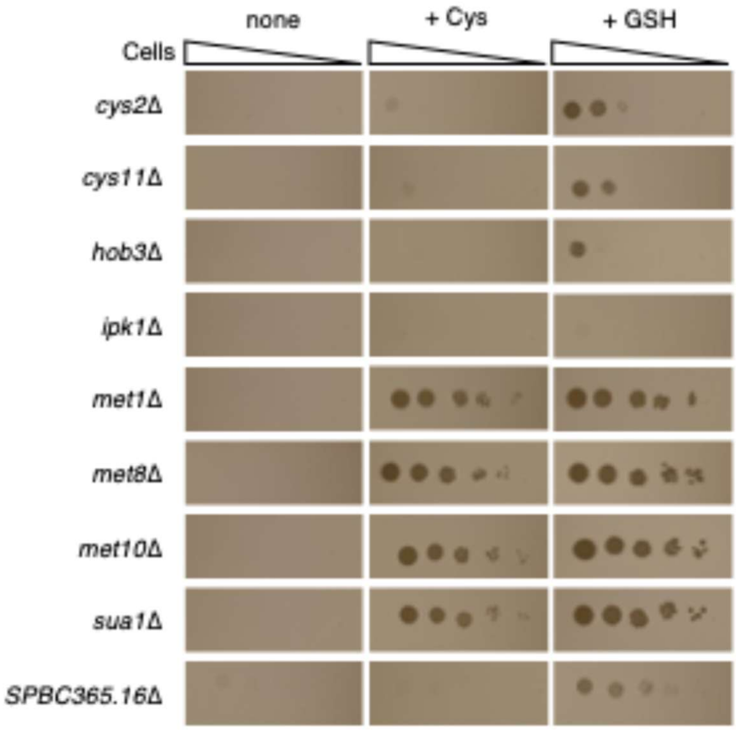


**S10 Figure. Growth recovery by cysteine or GSH on solid media.**
Serially diluted cells were spotted onto EMM2 (left), EMM2 supplemented with 2 mM Cys (center), or EMM2 supplemented with 3 mM GSH (right) and incubated at 27 °C for 7 days. *ipk1*Δ strain did not respond to GSH on agar medium. Representative images in 2 independent experiments are shown.

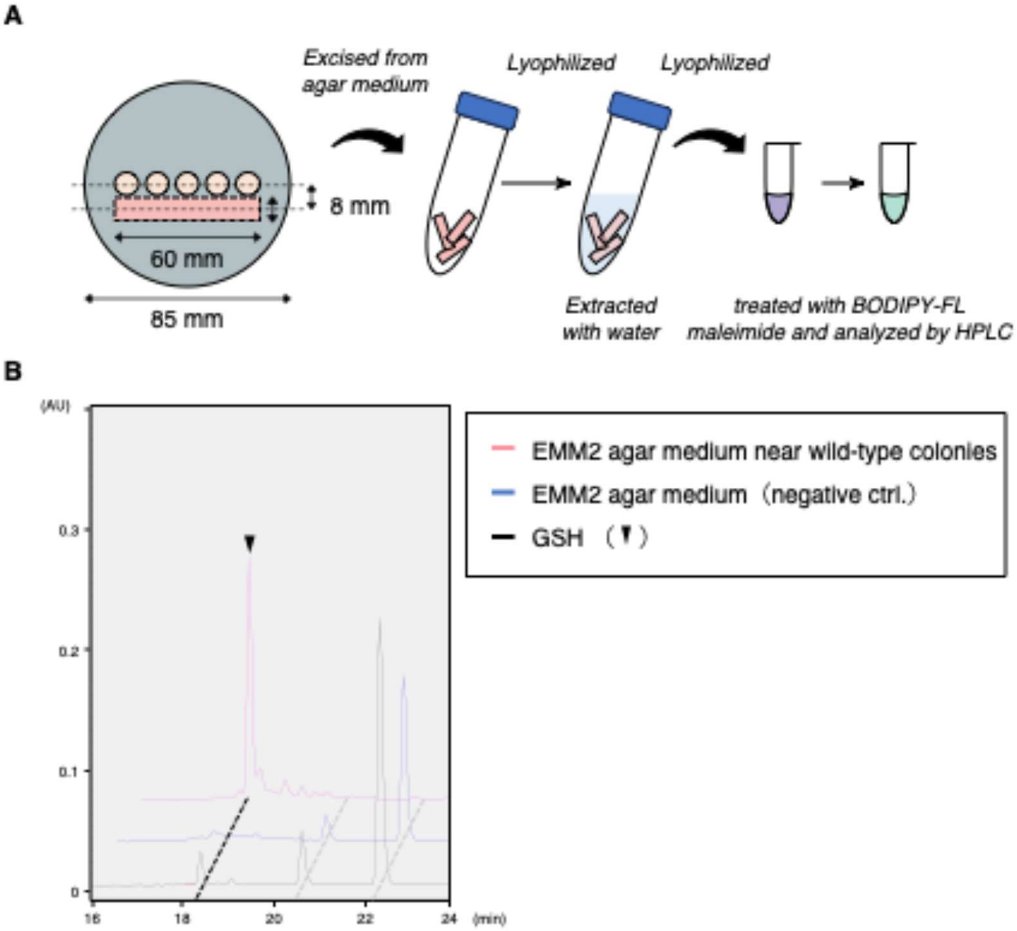


**S11 Figure. GSH was detected near the wild-type colonies on the agar medium.**

(A) Method for extracting metabolites from agar media. Wild-type colonies were cultured on EMM2 agar media at 27 °C for 7 days. Agar media surrounding the colonies were excised, lyophilized, and soaked in ultrapure water for 30 minutes. The extract was collected, lyophilized, and derivatized with BODIPY-FL maleimide followed by HPLC analyses.

(B) PDA analysis of EMM2 agar sample near wild-type colonies, EMM2 agar medium alone (negative control), and a GSH conjugate. Black arrowhead shows the GSH conjugate. The gray dotted lines indicate the BODIPY-FL maleimide alone. Representative images in 2 independent experiments are shown.

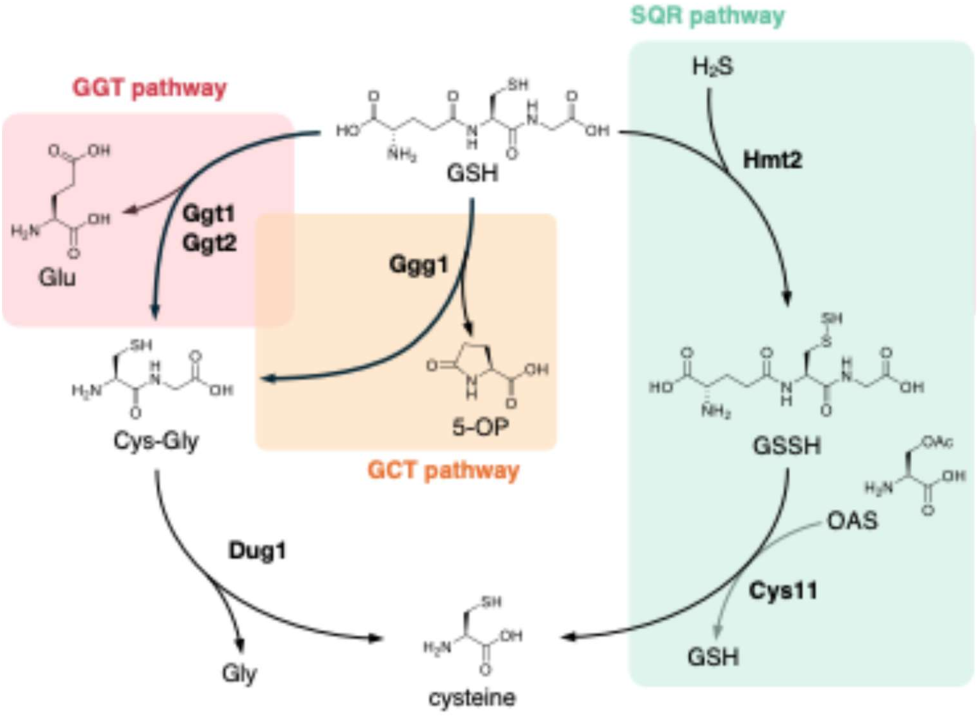


**S12 Figure. Assimilation pathways of GSH.**

Multiple pathways contribute to GSH assimilation. The γ-glutamyl transpeptidase (GGT) and γ-glutamyl cyclotransferase (GCT) pathways hydrolyze GSH to release cysteine via the intermediate Cys-Gly (left). In the sulfide quinone reductase (SQR) pathway, GSH is oxidized to GSSH, a sulfur donor for synthesizing Cys (right). In *S. pombe*, the double mutant lacking both Dug1 and Hmt2 shows a significantly reduced ability to assimilate GSH compared to either single mutant (S13 Fig), suggesting that both of the GGT/GCT and SQR pathways are functional.

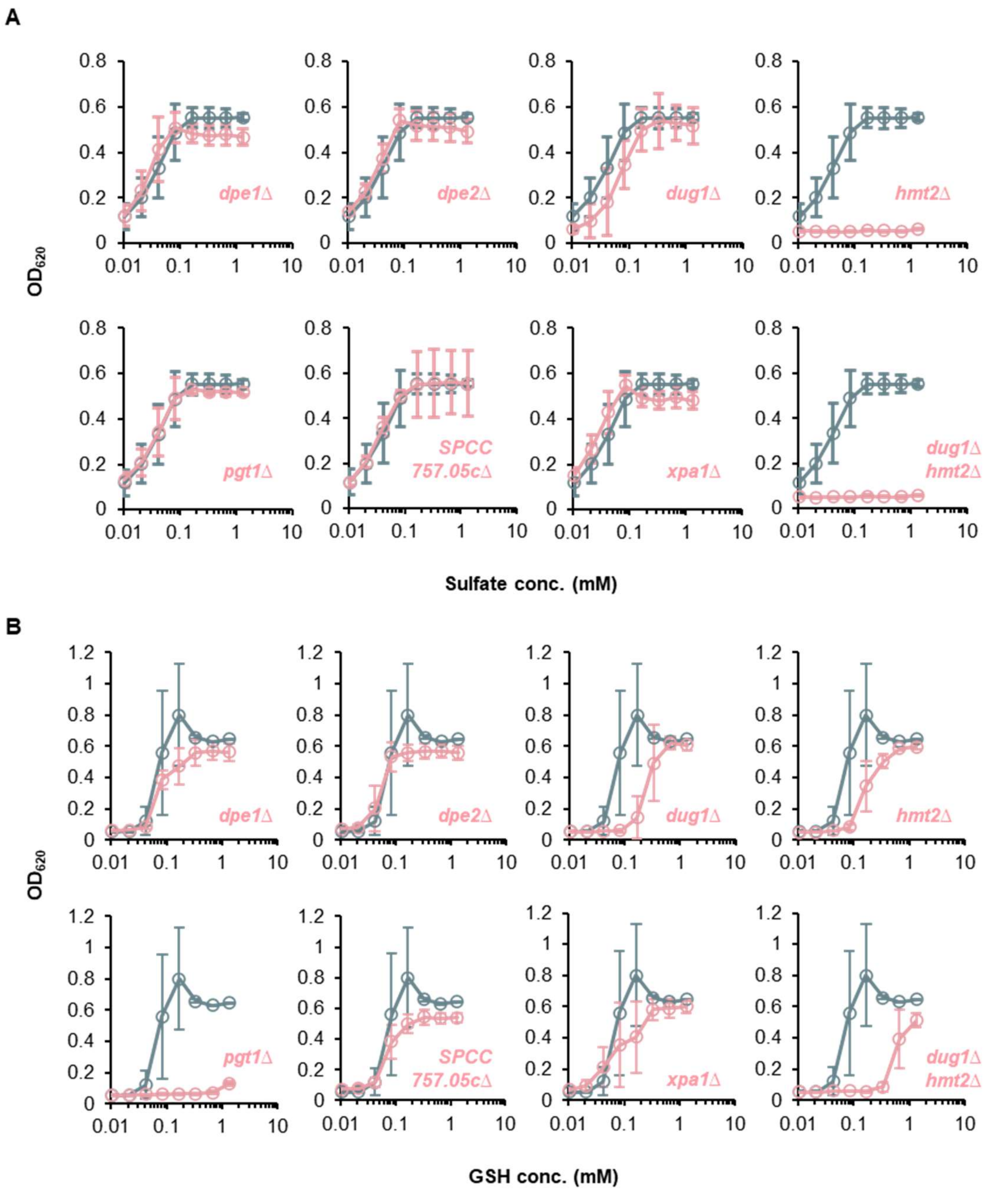


**S13 Figure. Sulfate and GSH assimilation as sulfur sources in gene deletion mutants.**

(A) Growth of mutants were measured in EMM2–S medium supplemented with the indicated concentrations of sulfate.

(B) Growth of mutants were measured in EMM2–S medium supplemented with the indicated concentrations of GSH.

Gray and red lines represent respectively wild-type and 8 mutants, respectively. Seven single-gene deletion mutants and one double-gene deletion mutant (*dug1*Δ *hmt2*Δ) were tested. Results for *dug1*Δ are also shown in Fig. 4A. The cultures were incubated at 30 °C for 48 hours. Data represent the mean ± SD (n=3).

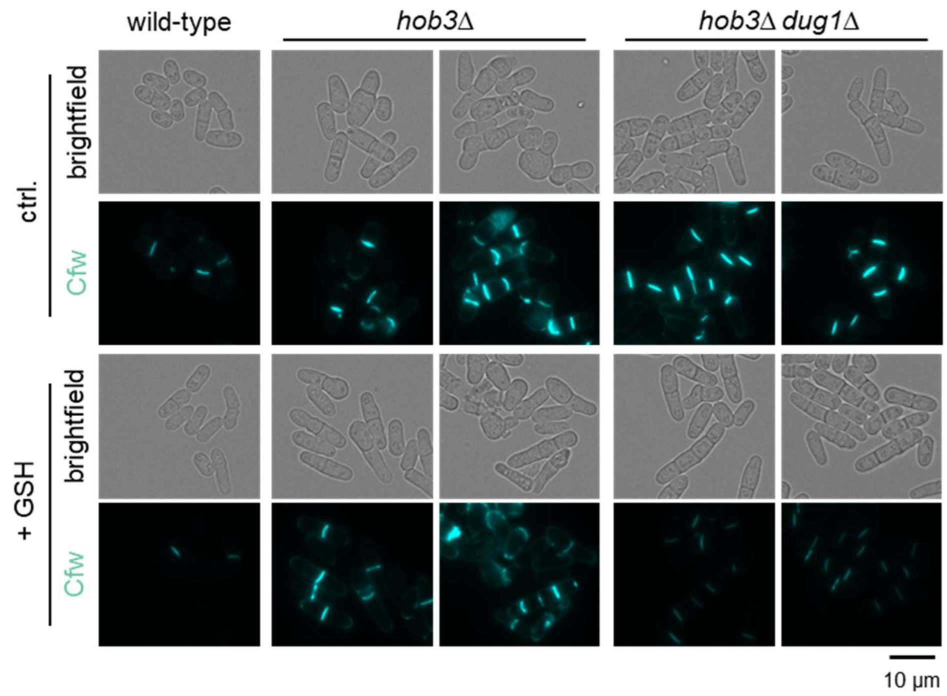


**S14 Figure. Morphological analyses of *hob3* mutants.**

Wild-type, *hob3*Δ, and *hob3*Δ *dug1*Δ cells were cultivated in EMM2 with or without 1 mM GSH and observed by fluorescence microscopy. Septa were visualized with calcofluor white (Cfw). Enlarged images are shown in Fig 5A. Scale bar = 10 μm. Representative images in 3 independent experiments are shown.

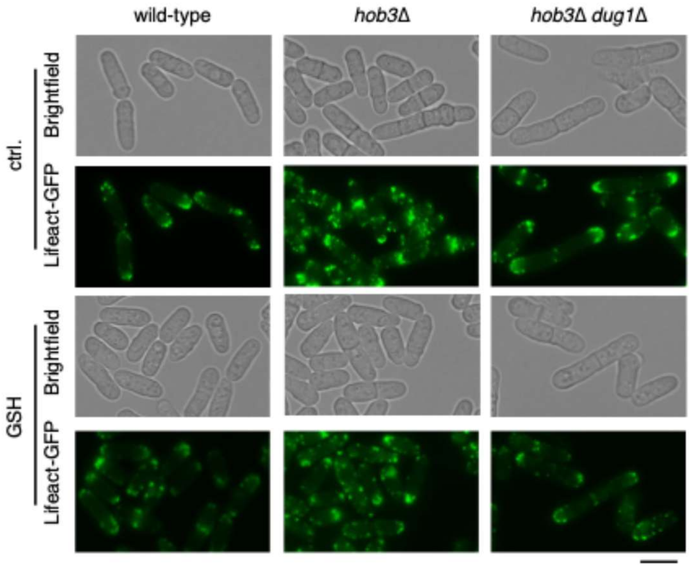


**S15 Figure. The observation of lifeact-GFP in *hob3* mutants.**

Wild-type, *hob3*Δ, and *hob3*Δ *dug1*Δ cells were cultivated in EMM2 with or without 1 mM GSH and observed by fluorescence microscopy. Scale bar = 10 μm. Representative images in 3 independent experiments are shown.

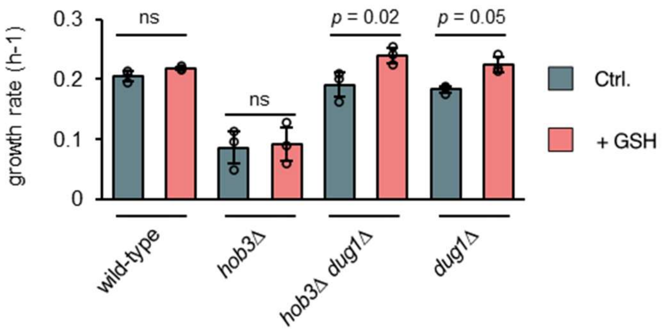


**S16 Figure. Growth speed of *hob3* mutants.**

Growth rates were calculated from $OD_{595}$ values obtained during the logarithmic growth phase. Cells were cultivated in EMM2 liquid media with or without 1 mM GSH at 30 °C. Data represent the mean ± SD (n = 3). P-values were determined using an unpaired two-tailed t-test, with Benjamini–Hochberg correction for multiple testing. ns, not significant.

S1 Table. List of 77 strains that showed growth on YES medium but no or poor growth on EMM2

This material is uploaded as an excel file.

S2 Table. GO enrichment analysis of 77 gene deletion mutants

| Fold Enrichment | Enrichment FDR | Pathway |
|---|---|---|
| 35.5 | 2.5E-07 | GO:0006526 arginine biosynthetic proc. |
| 27.4 | 8.9E-09 | GO:0009084 glutamine family amino acid biosynthetic proc. |
| 16.4 | 3.7E-26 | GO:0008652 cellular amino acid biosynthetic proc. |
| 16.3 | 1.2E-07 | GO:0000097 sulfur amino acid biosynthetic proc. |
| 16.1 | 2.0E-22 | GO:1901607 alpha-amino acid biosynthetic proc. |
| 13.0 | 7.3E-07 | GO:0000096 sulfur amino acid metabolic proc. |
| 12.1 | 2.9E-23 | GO:0046394 carboxylic acid biosynthetic proc. |
| 12.0 | 2.9E-23 | GO:0016053 organic acid biosynthetic proc. |
| 10.7 | 8.0E-07 | GO:0009066 aspartate family amino acid metabolic proc. |
| 10.5 | 9.0E-18 | GO:1901605 alpha-amino acid metabolic proc. |
| 10.4 | 2.5E-07 | GO:0044272 sulfur compound biosynthetic proc. |
| 9.0 | 5.8E-19 | GO:0006520 cellular amino acid metabolic proc. |
| 7.9 | 7.7E-21 | GO:0044283 small molecule biosynthetic proc. |
| 6.6 | 9.0E-18 | GO:0019752 carboxylic acid metabolic proc. |
| 6.4 | 1.6E-17 | GO:0043436 oxoacid metabolic proc. |
| 6.4 | 1.7E-17 | GO:0006082 organic acid metabolic proc. |
| 4.3 | 3.1E-16 | GO:0044281 small molecule metabolic proc. |
| 3.0 | 1.3E-11 | GO:1901566 organonitrogen compound biosynthetic proc. |
| 2.0 | 4.6E-07 | GO:1901576 organic substance biosynthetic proc. |
| 2.0 | 4.9E-07 | GO:0009058 biosynthetic proc. |

S3 Table. Summary of the growth recovery of 37 gene deletion mutants

This material is uploaded as an excel file.

S4 Table. GO enrichment analysis of 37 gene deletion mutants

| Fold Enrichment | Enrichment FDR | Pathway |
|---|---|---|
| 6.1 | 8.0E-03 | GO:0006091 generation of precursor metabolites and energy |
| 21.4 | 7.6E-03 | GO:0006534 cysteine metabolic process |
| 10.4 | 2.5E-03 | GO:0044272 sulfur compound biosynthetic process |
| 16.8 | 2.0E-03 | GO:0006555 methionine metabolic process |
| 35.7 | 1.8E-03 | GO:0006123 mitochondrial electron transport cytochrome c to oxygen |
| 35.7 | 1.8E-03 | GO:0019344 cysteine biosynthetic process |
| 9.0 | 1.2E-03 | GO:0015980 energy derivation by oxidation of organic compounds |
| 47.6 | 8.5E-04 | GO:0000103 sulfate assimilation |
| 22.0 | 8.5E-04 | GO:0008535 respiratory chain complex IV assembly |
| 22.0 | 8.5E-04 | GO:0033617 mitochondrial cytochrome c oxidase assembly |
| 15.9 | 4.8E-04 | GO:0000096 sulfur amino acid metabolic process |
| 11.4 | 4.4E-04 | GO:0045333 cellular respiration |
| 11.7 | 4.1E-04 | GO:0046034 ATP metabolic process |
| 12.6 | 2.9E-04 | GO:0009060 aerobic respiration |
| 18.3 | 2.9E-04 | GO:0033108 mitochondrial respiratory chain complex assembly |
| 19.8 | 2.4E-04 | GO:0000097 sulfur amino acid biosynthetic process |
| 20.4 | 2.3E-04 | GO:0017004 cytochrome complex assembly |
| 18.2 | 5.0E-05 | GO:0006119 oxidative phosphorylation |
| 57.2 | 3.7E-05 | GO:0006122 mitochondrial electron transport ubiquinol to cytochrome c |
| 23.8 | 1.4E-05 | GO:0022900 electron transport chain |
| 26.0 | 9.8E-06 | GO:0022904 respiratory electron transport chain |
| 30.6 | 4.5E-06 | GO:0019646 aerobic electron transport chain |
| 30.6 | 4.5E-06 | GO:0042773 ATP synthesis coupled electron transport |
| 30.6 | 4.5E-06 | GO:0042775 mitochondrial ATP synthesis coupled electron transport |

### S5 Table. Fission yeast strains used in this study.

| Strain | Genotype | Notes |
|---|---|---|
| JY1 | $h^{-}$ *wild-type* | Lab stock |
| MBY7200 | $h^{-}$ *pAct1-Lifeact-GFP::leu1 ade6-M216 ura4-D18 leu1-32* | Lab stock |
| RY74 | $h^{-}$ *dug1::hph*$^{r}$ *ade6-M216 ura4-D18 leu1-32* | This study |
| RY80 | $h^{?}$ *met10::kan*$^{r}$ *dug1::hph*$^{r}$ *ade6-M216 ura4-D18 leu1-32* | This study |
| RY83 | $h^{?}$ *SPBC365.16::kan*$^{r}$ *dug1::hph*$^{r}$ *ade6-M216 ura4-D18 leu1-32* | This study |
| RY86 | $h^{?}$ *met1::kan*$^{r}$ *dug1::hph*$^{r}$ *ade6-M216 ura4-D18 leu1-32* | This study |
| RY89 | $h^{?}$ *met8::kan*$^{r}$ *dug1::hph*$^{r}$ *ade6-M216 ura4-D18 leu1-32* | This study |
| RY95 | $h^{+}$ *hob3::kan*$^{r}$ *dug1::hph*$^{r}$ *ade6-M216 ura4-D18 leu1-32* | This study |
| RY98 | $h^{?}$ *cys11::kan*$^{r}$ *dug1::hph*$^{r}$ *ade6-M216 ura4-D18 leu1-32* | This study |
| RY101 | $h^{?}$ *hmt2::kan*$^{r}$ *dug1::hph*$^{r}$ *ade6-M216 ura4-D18 leu1-32* | This study |
| RY107 | $h^{?}$ *cys2::kan*$^{r}$ *dug1::hph*$^{r}$ *ade6-M216 ura4-D18 leu1-32* | This study |
| RY110 | $h^{?}$ *sua1::kan*$^{r}$ *dug1::hph*$^{r}$ *ade6-M216 ura4-D18 leu1-32* | This study |
| RY116 | $h^{-}$ *pgt1::hph*$^{r}$ *ade6-M216 ura4-D18 leu1-32* | This study |
| RY134 | $h^{+}$ *hob3::kan*$^{r}$ *pgt1::hph*$^{r}$ *ade6-M216 ura4-D18 leu1-32* | This study |
| RY152 | $h^{?}$ *hob3::kan*$^{r}$ *pAct1-Lifeact-GFP::leu1 ade6-M216 ura4-D18 leu1-32* | This study |
| RY155 | $h^{?}$ *hob3::kan*$^{r}$ *dug1::hph*$^{r}$ *pAct1-Lifeact-GFP::leu1 ade6-M216 ura4-D18 leu1-32* | This study |

### S6 Table. Oligo DNAs used in this study.

| Name | Sequence | Purpose |
|---|---|---|
| kan0 | GGTTAGGATTTGCCACTGAG | Confirmation of gene deletion |
| kan5 | TTTCTGCGCACTTAACTTCGC | |
| dug1_F1 | ATGGCGCAGTAAACCTATCATAGC | Gene deletion of dug1 |
| dug1_F2 | CCTGTTATCCCTAGCGGATCTGACGTTCTTCTGCGCTGCTAG | |
| dug1_R1 | TTCGCAGGTGTTGCTGGAAG | |
| dug1_R2 | GAGAGGCGGTTTGCGTATTGTGCGAGCCTTGGTATGTAACTG | |
| dug1_checkF | TGAATTAAGGCCAGCTCCTGTC | Confirmation of gene deletion |
| dug1_checkR | ACTTCGCCGTTCTACCATTGAC | |
| pgt1_F1 | ATGGTTCTGATGCTGCTTTG | Gene deletion of pgt1 |
| pgt1_F2 | CCTGTTATCCCTAGCGGATCTGGTGTCCCTTCGCTCTTTATCG | |
| pgt1_R1 | CTGATCCAGCACAGAATGTTCC | |
| pgt1_R2 | GAGAGGCGGTTTGCGTATTGAGGGCTTGCCTGTCCATATAC | |
| pgt1_check_F | ACGAGGTCAAGTACCTCTTCA | Confirmation of gene deletion |
| pgt1_check_R | ACTCTGACTGTCCGTTCATCTC | |
